# SARS-CoV-2 nucleocapsid protein inhibits the PKR-mediated integrated stress response through RNA-binding domain N2b

**DOI:** 10.1101/2022.09.02.506332

**Authors:** Chiara Aloise, Jelle G. Schipper, Arno van Vliet, Judith Oymans, Tim Donselaar, Daniel L. Hurdiss, Raoul J. de Groot, Frank J.M. van Kuppeveld

## Abstract

The nucleocapsid protein N of severe acute respiratory syndrome coronavirus 2 (SARS-CoV-2) enwraps and condenses the viral genome for packaging but is also an antagonist of the innate antiviral defense. It suppresses the integrated stress response (ISR), purportedly by interacting with stress granule (SG) assembly factors G3BP1 and 2, and inhibits type I interferon responses. To elucidate its mode of action, we systematically deleted and over-expressed distinct regions and domains. We show that N via domain N2b blocks PKR-mediated ISR activation, as measured by suppression of ISR-induced translational arrest and SG formation. N2b mutations that prevent dsRNA binding abrogate these activities also when introduced in the intact N protein. Substitutions reported to block post-translation modifications of N or its interaction with G3BP1/2 did not have a detectable additive effect. In an encephalomyocarditis virus-based infection model, N2b - but not a derivative defective in RNA binding - prevented PKR activation, inhibited β-interferon expression and promoted virus replication. Apparently, SARS-CoV-2 N inhibits innate immunity by sequestering dsRNA to prevent activation of PKR and RIG-I-like receptors. Similar observations were made for the N protein of human coronavirus 229E, suggesting that this may be a general trait conserved among members of other orthocoronavirus (sub)genera.

**SIGNIFICANCE STATEMENT:** SARS-CoV-2 nucleocapsid protein N is an antagonist of innate immunity but how it averts virus detection by intracellular sensors remains subject to debate. We provide evidence that SARS-CoV-2 N, by sequestering dsRNA through domain N2b, prevents PKR-mediated activation of the integrated stress response as well as detection by RIG-I-like receptors and ensuing type I interferon expression. This function, conserved in human coronavirus 229E, is not affected by mutations that prevent posttranslational modifications, previously implicated in immune evasion, or that target its binding to stress granule scaffold proteins. Our findings further our understanding of how SARS-CoV-2 evades innate immunity, how this may drive viral evolution and why increased N expression may have been a selective advantage to SARS-CoV-2 variants of concern.

## INTRODUCTION

Vertebrate cells are provided with a diversity of interconnected sensors and effectors to timely detect and counter viral infection. Particularly dsRNA, an inevitable product of RNA and DNA virus replication, triggers a vigorous intracellular antiviral response (1). For example, binding of dsRNA by 2’-5’oligoadenylate (2-5A) synthase (OAS) leads to enzyme activation and production of 2-5A, which in turn activates RNase L to stall the synthesis of viral proteins through non-specific RNA degradation (2-7). Another dsRNA-activated pathway, the integrated stress response (ISR) with proteinase kinase R (PKR) as key sensor, entails inhibition of translation initiation and ultimately cell death. Upon dsRNA binding, PKR dimerizes, autophosphorylates and then proceeds to phosphorylate the alpha subunit of eukaryotic initiation factor eIF2, turning it into competitive inhibitor of guanine nucleotide exchange factor eIF2b. In consequence, the production of eIF2-GTP-(Met)tRNA_i_^Met^ ternary complex is downregulated, recognition of the initiation codon is blocked and cap-dependent translation-initiation prevented (8-11). Polysome dissociation, ensuing activation of the stress response, results in an excess of stalled 48S preinitiation complexes, which are stored in cytoplasmic membraneless organelles called stress granules (SGs) (12-14).

SGs are dynamic ribonucleoprotein assemblies that are formed through liquid-liquid phase separation with RNA binding protein Ras GTPase-activating protein-binding proteins 1 and 2 (G3BP1 and -2) functioning both as a molecular switches and main protein scaffolds together with T-cell-restricted intracellular antigen 1 (TIA1) and TIA1-related protein (TIAR) (15-19). The SGs thus serve as deposits from which mRNAs, poised for translation through association with critical components of the translation machinery (40S ribosomal subunits, eIF4E, eIF4G, eIF4A, eIF4B, Poly(A) binding protein, eIF3, and eIF2), can be rapidly retrieved. SGs, however, are also considered a coordinating hub for the activation of other antiviral defense mechanisms like those of RIG-I-like receptors (RLRs). Indeed, RIG-I and MDA5, RLR-regulating PKR-activating protein (PACT), RLR-modulating ubiquitin ligases TRIM25 and TRAF2, and polyubiquitin chains are all recruited to ISR-induced SGs (20-23).

The antiviral mechanisms elicited by dsRNA are highly effective, even to such an extent that all known mammalian viruses code for one or more antagonists (11). Coronaviruses, positive-stranded RNA viruses of exceptional genetic complexity, code for a universally conserved endonuclease (EndoU) that efficiently prevents simultaneous activation of host cell dsRNA sensors OAS, PKR and MDA5 through dsRNA decay (24-26). Illustrating the importance of EndoU, mutants defective for EndoU are severely attenuated *in vivo*, and their replication in cultured primary macrophages is restricted presumably due to high basal expression levels of host sensors in these cells (25). EndoU is derived by proteolytic processing of a large replicase polyprotein pp1ab, translated from the incoming viral genome, and essential to evade early innate and intrinsic antiviral host cell responses.

Apparently, however, EndoU may not be sufficient to suppress dsRNA-mediated antiviral activities during later stages of the viral life cycle. To express the open reading frames (ORFs) downstream of the replicase gene, CoVs produce a 3’ co-terminal nested set of subgenomic mRNAs from which the structural proteins are translated in addition to a variety of so-called accessory non-structural proteins. Some of the latter also have been shown to counteract dsRNA-mediated antiviral host responses. For example, the ns4a protein of Middle East Respiratory Syndrome coronavirus (MERS-CoV) prevents PKR-mediated stress by sequestering dsRNA (27, 28), whereas the MERS-CoV ns4b protein is a phosphodiesterase (PDE) and antagonizes the OAS-RNase L pathway by enzymatically degrading 2-5A activators (29). Non-related PDEs, ns2 proteins, are found in members of the subgenus *Embecovirus*, including human coronavirus OC43 (30). Finally, a specific inhibitor of the ISR was found in gammacoronaviruses of cetaceans. The beluga whale coronavirus accessory protein 10 (BWCoV acP10) blocks p-eIF2-eIF2B association to allow continued formation of the ternary complex and unabated global translation even at high p-eIF2 levels that would otherwise cause translational arrest (31).

Apparently, CoVs rely on redundancy in antagonists and antagonistic mechanisms to effectively counter dsRNA-induced antiviral host responses. Likewise, the current pandemic severe acute respiratory syndrome coronavirus 2 (SARS-CoV-2) codes for an ISR inhibitor additional to EndoU (32). Its nucleocapsid protein (N) was reported to block the ISR and to inhibit SG formation in a PKR- and G3BP1-dependent fashion (33-36). In apparent concordance, proteomics studies and structural analyses provided evidence for physical interactions between N and the SG key components G3BP1 and G3BP2 (33, 37-40).

Coronavirus N proteins have a modular structure with N-terminal and C-terminal RNA-binding domains, called N1b and N2b respectively, bounded by largely disordered regions. Here, we took a reductionistic approach involving systematic deletion and individual transient expression of the different regions and domains. We show that the N2b domain is critical and sufficient to counter the ISR. Single site mutations in N2b that block dsRNA binding prevent PKR activation. This activity is not affected by posttranslational modifications elsewhere in the N protein that are known to regulate RNA binding nor dependent on physical interaction with G3BP1 and G2BP2. Using the encephalomyocarditis virus as a surrogate infection system we show that N2b domain prevents PKR-mediated activation of the ISR and suppresses IFNβ expression also in virus-infected cells. Our findings suggest that in addition to a function in replication and genome packaging, SARS-CoV-2 N functions as a classical antagonist of dsRNA-induced host defense.

## RESULTS

### SARS-CoV-2 N inhibits PKR-induced ISR

Transfection of cells with specific expression plasmids like pEGFP-N3 triggers the ISR through dsRNA-mediated PKR activation, resulting in translation arrest and the formation of SGs (**Fig. 1A**) (27, 41-43). This phenomenon allows for a convenient method to identify potential viral ISR antagonists by transiently expressing these proteins genetically fused to enhanced green fluorescent protein (EGFP) (27, 31). The expression levels of these fusion proteins, as judged by fluorescence microscopy, can then be compared to that of EGFP alone as an indicator for translation arrest and the prevention thereof. In addition, SG formation can be assessed by immunofluorescence analysis (IFA) by staining the transfected cells for established SG markers like G3BPs or eIF3. This procedure previously allowed us to identify several stress antagonists including MERS-CoV 4a and Beluga whale coronavirus AcP10 (27, 31).

**Fig. 1.**
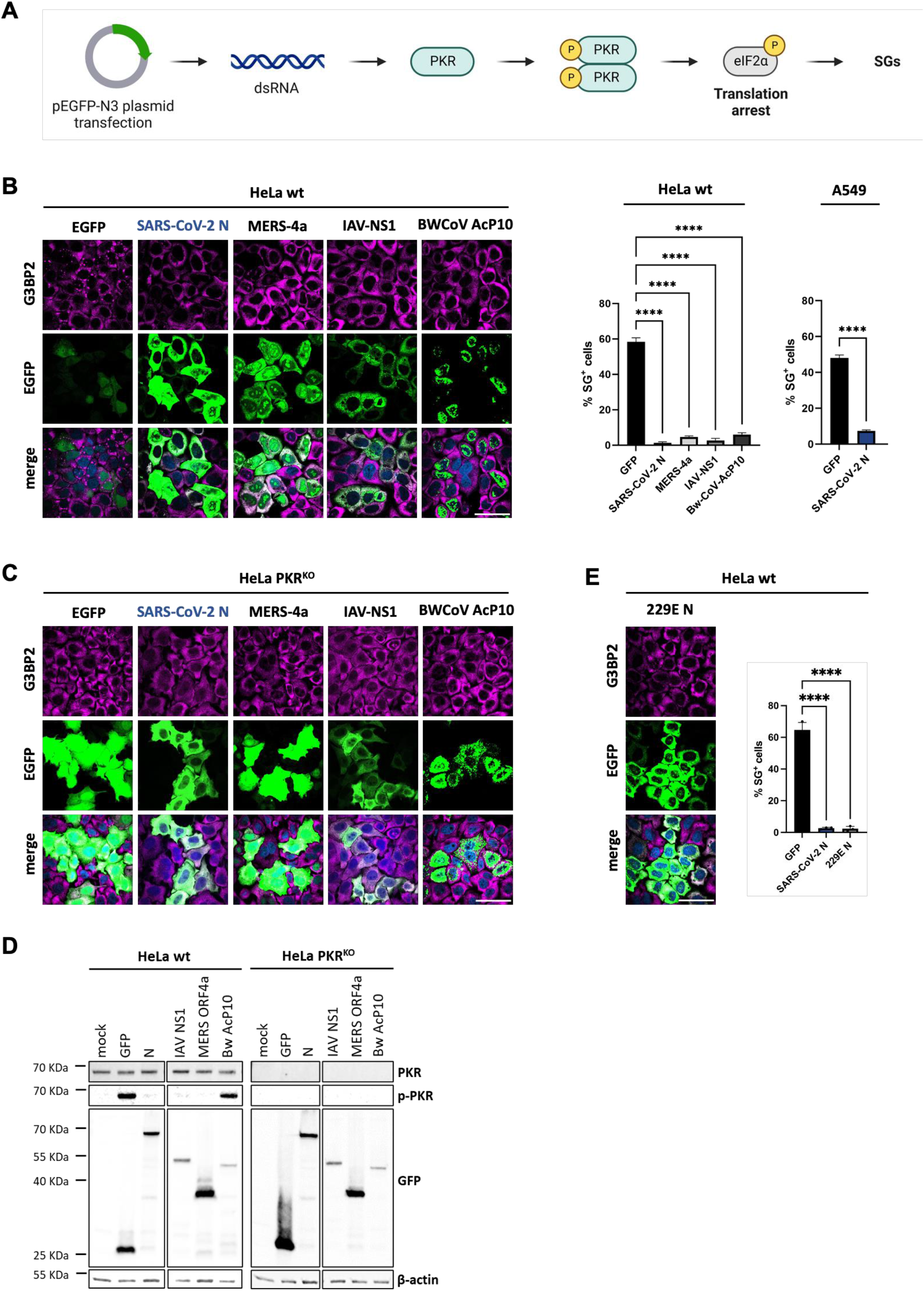
SARS-CoV-2 N inhibits PKR-induced ISR, preventing translation arrest and SG formation in HeLa and A549 cells. (***A***) Schematic representation of the PKR-induced ISR activation upon the pEGFP-N3 transfection in eukaryotic cells. Created with BioRender.com. HeLa and A549 cells were transfected with expression vector pEGFP-N3 (EGFP) or pEGFP-N3 derivatives encoding SARS-CoV-2 N, MERS-4a, IAV-NS1 or Bw-CoV-AcP10 genetically fused to EGFP. Induction of the ISR via dsRNA-mediated PKR activation or suppression thereof was assessed by comparing EGFP fluorescence intensity and SG formation as detected by immunofluorescence staining for G3BP2. (left panel) Transfected HeLa cells; (immune)fluorescence microscopy images, representative results. Scale bars: 50 µm. (right panel) Bar graphs for Hela and A549 cells showing the percentage of EGFP expressing cells containing G3BP2-positive SGs. The results are representative of three independent experiments counting >200 cells per sample. Standard deviation indicated by error bars; *** p=0.001, **** p< 0.001, ns = not significant (one-way ANOVA with Dunnett post hoc test). HeLa PKR^KO^ cells were transfected as in B. Representative (immuno)fluorescence microscopy images are shown. (***D***) Western-blot analysis for PKR, phosphorylated PKR (p-PKR), GFP-fused proteins and β-actin. HeLa wt and PKR^KO^ cells, mock-transfected or transfected with indicated plasmid were lysed at 24 hr. Of note, the larger yield of EGFP versus EGFP-N is counter intuitive but can be explained from the fact that (i) this is an ensemble measurement (for the total transfected cell population) versus the analysis of individual cells by fluorescence microscopy and (ii) basal expression levels of the codon optimized EGFP *prior* to ISR activation will exceed those the EGFP-N fusion protein, which is non-codon optimized and three times larger in size (see also **Fig. S2**). (***E***) Inhibition of SG formation and translational arrest by pEGFP-N3-expressed N proteins of MERS-CoV and HCoV-229E. (Immuno)fluorescence analysis (left panel), quantitative representation of the results and statistical analysis as in B.

SARS-CoV-2-infected Vero E6-TMPRRS2 cells, identified by detection of dsRNA, were virtually devoid of SGs, suggestive of virus-induced suppression of the ISR (**Fig. S1**). While probing SARS-CoV-2 proteins for a potential role in ISR inhibition, we noted that in transfected wildtype (wt) HeLa cells, the expression levels of the SARS-CoV-2 N-EGFP fusion protein were strongly increased as compared to the EGFP control. This pattern of enhanced expression in a sizeable population of transfected cells was similar to that observed for established ISR antagonists (**Fig. 1B**). Moreover, like these antagonists, SARS-CoV-2 N significantly reduced the number of transfected cells with stress granules both in wildtype HeLa (HeLa wt) as well as in A549 cultures (**Fig. 1B**). SG suppression was observed also in cells with fluorescence intensities just above the background, suggesting that low levels of N-EGFP are already sufficient to prevent SGs to form.

To confirm that expression of EGFP was suppressed due to PKR-induced ISR-mediated translational arrest, we repeated the experiments in PKR-deficient HeLa PKR^KO^ cells. Indeed, EGFP expression was increased and SGs were absent in these cells (**Fig. 1C**). In further accordance, PKR was activated -as indicated by PKR phosphorylation-in HeLa wt cells expressing EGFP but not in cells expressing EGFP-N. (**Fig. 1D**, left panel). Finally, Western blot analysis confirmed that EGFP-N was expressed to similar levels in HeLa wt and PKR^KO^ cells. Levels of EGFP, however, were strongly increased (about 7-fold) in HeLa PKR^KO^ cells (**Fig. 1D**, right panel). The combined findings identify SARS-CoV-2 N as an ISR antagonist that prevents translational arrest and ensuing SG formation by acting as a PKR inhibitor.

The N protein of another betacoronavirus, MERS-CoV, was previously noted to suppress SG formation too (35). Interestingly, we observed inhibition of translational arrest and SG formation also in cells expressing the N protein of human alphacoronavirus 229E (HCoV-229E) (**Fig. 1E**). Apparently, N-mediated suppression of the ISR is not unique to betacoronaviruses but may well be a general trait conserved among members of other orthocoronavirus (sub)genera.

### Domain N2b is essential and sufficient for suppression of PKR-induced SG formation

The N proteins from the four CoV genera share only 27 to 30% sequence identity but are conserved in their modular organization (**Fig. 2A**) ((44, 45) for a review see (46)). They are comprised of two ordered domains, the N-terminal domain (N1b also called NTD) and the C-terminal domain (N2b aka CTD) (in SARS-CoV-2 N, residues 49-175 and 248-365, respectively). N1b and N2b are separated and flanked by segments, predicted to be at least partially disordered (N1a, N2a and N3). N1b and N2b have been implicated in RNA binding and, in case of N2b, also in N dimerization. Indeed, in infected cells and upon heterologous expression, CoV N proteins form homodimers that, in turn, assemble into tetramers mediated by N3 (47). The N2a central spacer contains a serine- and arginine-rich (SR) region, immediately downstream of N1b, the regulated phosphorylation of which is deemed important for N function during different stages of the CoV replication cycle (48-50). The primary function of N is in virus assembly. It condenses newly produced gRNA into helical nucleocapsids, apparently with N2b controlling target selectivity (51, 52), and then drives envelopment by binding to the viral membrane protein M via N3. However, N has several functions auxiliary to virion morphogenesis. At the very start of the infectious cycle, N is essential for the initiation of infection by the incoming gRNA through association of SR with the cytosol-exposed ubiquitin1 domain of replication organelle pore protein nsP3 (50, 53). Moreover, purportedly relevant to ISR suppression, N binds to G3BP1 and 2 through segment N1a (38, 40, 54).

**Fig. 2.**
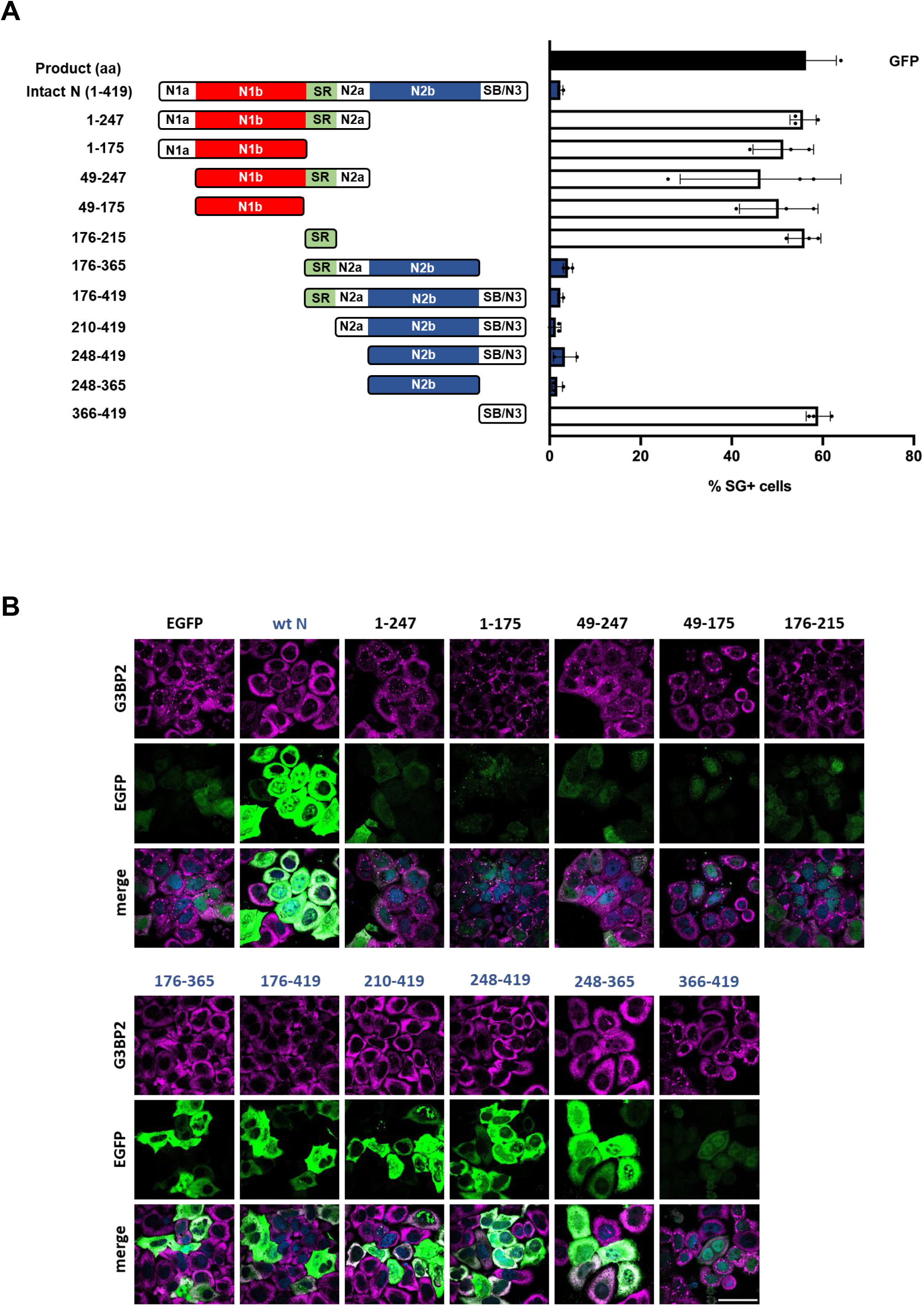
Nucleocapsid domain N2b suppresses translational arrest and SG formation. (***A***) Schematic representation of truncated derivatives of SARS-CoV-2 N protein fused to EGFP (left panel). The proteins were overexpressed from pEGFP-N3-based vectors in HeLa cells and percentages of SG-positive cells were determined as in Fig. 1A (right panel). The data are representative of three independent experiments with more than 200 cells counted for the presence of G3BP2-positive SGs per individual sample. Standard deviation indicated by error bars. For a statistical analysis of the results, see **Table S1**. (***B***) Representative results from (immuno)fluorescence analysis of HeLa cells transfected to express the SARS-CoV-2 N protein derivatives. Scale bar: 50 µm.

To determine the molecular basis for ISR suppression, we constructed a library of N mutants with partially disordered regions and domains either systematically deleted or expressed in isolation. As shown in **Fig. 2**, suppression was lost upon deletion of subdomain N2b. Moreover, expression of N2b in isolation caused a reduction in SGs to an extent similar to that of full-length N. The data therefore suggested that ISR inhibition and SG suppression as observed for the intact N protein is mediated at least in part by N2b.

### N2b mutations that disrupt dsRNA binding abrogate suppression of SG formation

N2b binds both single and double stranded oligonucleotides, whether RNA or DNA (46, 55, 56). Hence, a possible mechanism for N2b to prevent PKR-induced activation of the ISR is by sequestering dsRNA. Crystal structural analysis revealed that N2b homodimers form a rectangular slab with wide faces of 45 Å × 35 Å in dimension (55). One is comprised of an interlaced inter-molecular four-stranded β-sheet and predominantly negatively charged, the other of two α-helical regions separated by a shallow central, positively charged groove thought to be the oligonucleotide binding site (46, 55, 57, 58). Within the groove, there are several conserved positively charged residues with their surface-exposed side chains seemingly poised for interaction with nucleic acid, in particularly Lys^257^ and Lys^261^ (**Fig. 3A**). The binding by bacterially-expressed N2b of synthetic RNA oligonucleotides, whether single or double-stranded, SARS-CoV-2 derived (highly conserved 3’ UTR stem-loop II motif, s2m; (59)) or a scrambled version thereof, can be readily demonstrated by electrophoretic mobility assay (EMSA) (**Fig. 3A, B**).

**Fig. 3.**
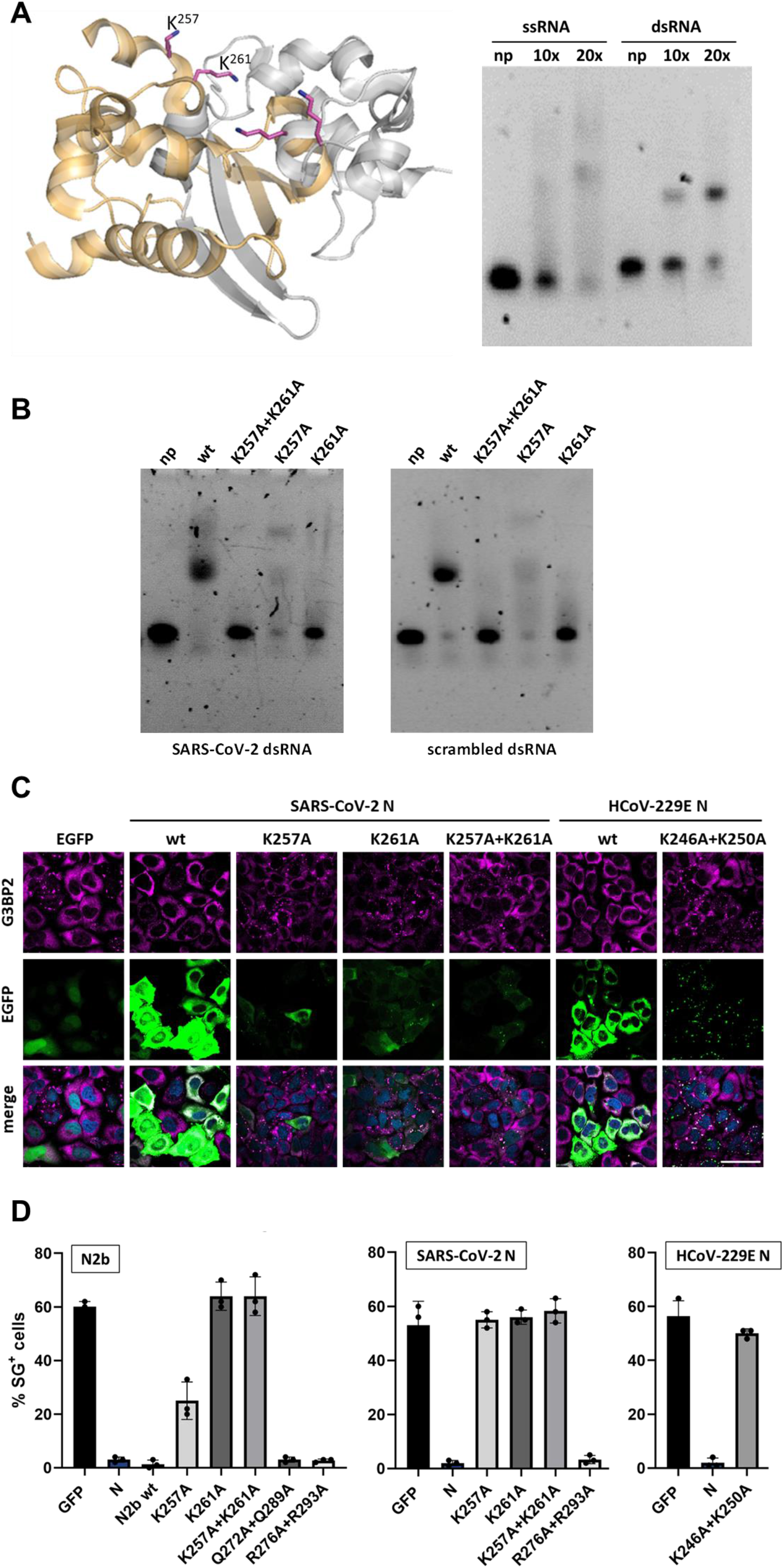
N2b mutations that disrupt dsRNA binding abrogate suppression of translational arrest and SG formation. (***A***) (left panel) Cartoon representation of the SARS-CoV-2 N2b dimer (PDB ID code 7C22 (58); generated with PYMOL with monomers in grey and orange. Top view displaying the positively charged central RNA-binding groove. Side chains of Lys^257^ and Lys^261^, shown for both monomers and marked for one, in sticks and colored by element with carbon atoms in magenta. (right panel) Electrophoretic mobility shift assay (EMSA) with bacterially expressed N2b and single (ss) and double-stranded (ds) RNA oligonucleotides, designed after SARS-CoV-2 stem-loop II motif (s2m). Assays were performed with N2b in 10- or 20-fold molar excess. Non-bound RNA was included as a control (np, no protein). (***B***) The effect of N2b amino acid substitutions on binding of s2m dsRNA (left) or a scrambled version thereof (right). EMSAs performed with N2b and mutant derivatives in 20-fold molar excess. (***C****-****D***) Mutational analysis of SARS-CoV-2 N2b, full-length SARS-CoV-2 N and full-length HCoV-229E N. Select surface-exposed charged residues were substituted by Ala either individually or in combination as indicated and the effect on IRS-induced translational arrest (C) and SG formation (D) was analyzed in HeLa cells as in Fig. 1. For a statistical analysis of the results, see **Table S2**.

To test whether Lys^257^ and Lys^261^ are involved in RNA binding and whether such binding is important to counteract the ISR and suppress SG formation, they were replaced by Ala, separately and in combination. Indeed, dsRNA binding was either significantly reduced or lost beyond detection upon substitution of Lys^257^ and Lys^261^, respectively (**Fig. 3B**). In accordance, upon Ala substitution of Lys^261^ or Lys^257^, the mutant N2b proteins no longer rescued protein synthesis (**Fig. 3C**) and lost their capacity to suppress the formation of SGs, either completely (Lys^261^) or partially (Lys^257^) (**Fig. 3D**). Notably, Ala substitution of other surface-exposed charged residues (Gln^272^, Gln^289^, Arg^276^, Arg^293^), not located within the RNA binding groove but rather implicated in N2b inter-dimer association (46, 56), did not detectably affect SG formation (**Fig. 3D**).

Importantly, also in the intact N protein, Lys^261^Ala substitution abrogated inhibition of translational arrest as well as SG formation and so did the Lys^257^Ala mutation (**Fig. 3C, D**). It is unknown why the latter mutation exerts a stronger effect in the context of the full-length protein than in N2b. The combined data, however, do show that Lys^261^ and Lys^257^ are required for nucleic acid binding by N2b and suggest that this capacity to bind RNA is ssential for SG suppression. Moreover, the observation that also for the intact N protein SG suppression was reduced to background levels by single site Lys^261^Ala and Lys^257^Ala substitutions suggests that inhibition of PKR-induced ISR by SARS-CoV-2 N critically relies on N2b and N2b-mediated RNA binding. Notably, the N protein of alphacoronavirus HCoV-229E also lost its capacity to block SG formation when the orthologous N2b residues, Lys^246^ and Lys^250^ were substituted (**Fig. 3C, D**), suggestive of a common mechanism for ISR inhibition through the binding of RNA.

### N2b-mediated inhibition of PKR-induced SG formation is not affected by posttranslational modifications of N

Previous studies identified N as a multifunctional protein involved in different aspects of the viral replication cycle beyond viral assembly. Its activity apparently is regulated by posttranscriptional modifications such as differential phosphorylation of the SR domain, which alters RNA binding affinity (60, 61), and acetylation at Lys^375^, reportedly essential for liquid–liquid phase separation of N-RNA ribonucleoprotein complexes (62). Arginine methylation of SARS-CoV-2 nucleocapsid protein at Arg^95^ and Arg^177^ was reported to regulate RNA binding and its ability to suppress stress granule formation (33).

To test the importance of these posttranscriptional modifications on N2b-mediated inhibition of SG formation, we performed site-directed mutagenesis in the context of the intact N protein. Suppression of SG formation was not affected by Ala substitution of Arg^95^or Lys^375^ indicating that methylation and acetylation of N is not essential (**Fig. 4A, B**).

**Fig. 4.**
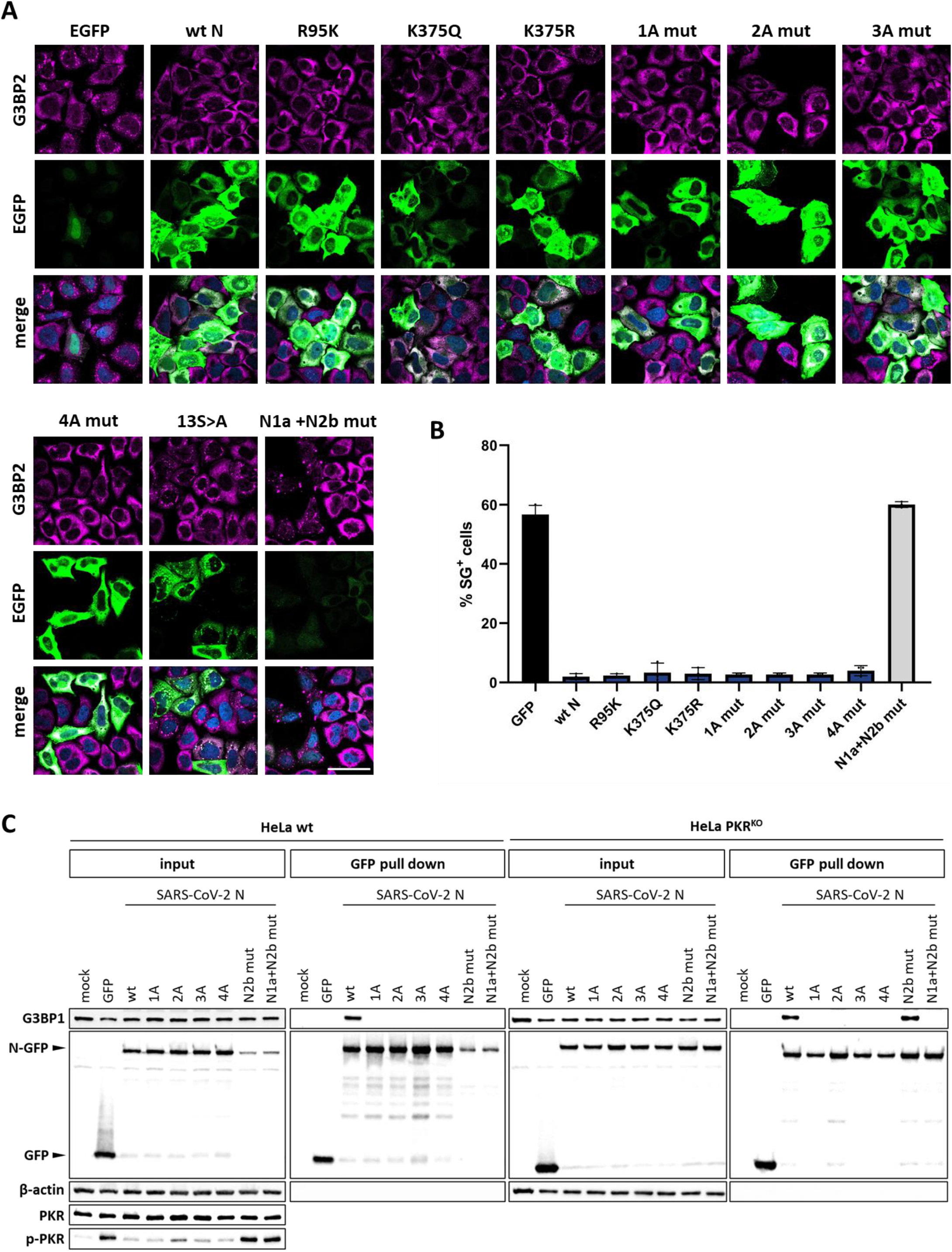
N-mediated suppression of SG formation and ISR-induced translational arrest is not affected by posttranslational modifications or G3BP1 interaction. (***A***) The effect on N-mediated ISR inhibition by mutations introduced to prevent posttranslational modifications (K^375^Q, K^375^R, R^95^K), to disrupt the G3BP1/2 binding site (1A-4A mutants) or to prevent phosphorylation of the SR element (13S>A) was tested in HeLa cells by (immune)fluorescence analysis as in Fig. 1. Scale bar: 50 µm. (***B***). Quantification of the results in (A) representative of three independent experiments, with >200 cells counted per sample as in Fig. 1. (***C***) Pull down assay to test SARS-CoV-2 N mutants for their association with endogenous G3BP1 in HeLa wt (left panel) and PKR^KO^ cells (right panel).

An N derivative in which phosphorylation of the SR segment was prevented through Ala substitution of 13 out of 14 Ser residues (13S>A) displayed a cellular distribution distinctively different from that of the parental wildtype protein, accumulating in large aggregates enriched for G3BP1 but mostly devoid of eIF3 (**Fig. 4A**; **Fig. S3**). These deposits differed in size, appearance and composition from typical SG. Importantly, the expression levels of the N-13S>A mutant were like those of wildtype N, as judged by fluorescence intensity observed in IFA, and consistently higher than those of EGFP (**Fig. 4A**). We interpret the findings to indicate that the SR mutations may cause the N protein to aggregate but do not affect inhibition of ISR-induced translational arrest. The results confirm those of our systematic deletion analysis (Fig. 2).

### N2b-mediated inhibition of SG formation is not affected by disruption of the G3BP binding motif

A ΦxFG motif within N1a (residues 15-18), required for association with G3BP1 and 2 (40), was recently proposed to rewire the G3BP interactome to disrupt stress granules (38). To investigate a possible role of N-G3BP interaction in suppressing PKR-induced SGs, we mutated the N1a ΦxFG motif through Ala substitution of key residue Phe^17^ (mut 1A) or by a combination of Ala substitions: Arg^14^ and Ile^15^ (mut 2A), Ile^15^, Phe^17^ and Gly^18^ (mut 3A), or Ile^15^, Thr^16^, Phe^17^ and Gly^18^ (mut 4A). In each case, binding to endogenous G3BP1 was no longer detectable by pull down assay (**Fig. 4C**) yet suppression of SG formation was not affected (**Fig. 4A, B**). Conversely, N proteins with N2b mutations to abrogate dsRNA binding no longer blocked the ISR (**Figs. 3** and **4B, C**) nor inhibited SG formation, even though N1a was still intact to bind G3BP1 to similar extent as wildtype N (**Fig. 4C**). Note that for these N2b mutants N-G3BP association could not readily be assessed in HeLa wt cells. As a result of PKR-activation (**Fig. 4C**) and ISR-mediated translational arrest, their expression levels were strongly reduced as compared to those of wt N and the amounts of coprecipitated G3BP1 were below the detection level of our Western blot analysis. Unperturbed G3BP binding, however, was evident, when assessed in HeLa PKR^KO^ cells (lane marked ‘N2b mut’) and lost upon concomitant mutation of the N1a ΦxFG G3BP binding motif (lane marked ‘N1a+N2b mut’) (**Fig. 4C**). The findings indicate that SARS-CoV-2 N-G3BP interaction is not required for inhibition of the ISR, at least not upon induction of the ISR via the PKR signaling pathway under the conditions applied. Moreover, this interaction is also not sufficient to prevent PKR-induced formation of SGs or to promote their disassembly.

To study whether G3BP-binding might still affect SG formation, HeLa PKR^KO^ cells were transfected to express wildtype N or mutant derivatives and subjected to arsenite/heme-regulated inhibitor kinase (HRI)-induced stress. This approach allowed us to study the importance of N-G3BP1 interaction more directly, i.e. without transfection induced PKR activation and N-mediated inhibition of PKR as complicating factors. Under these conditions, N inhibited SG formation albeit to limited extent (**Fig. 5**). The observed reduction in the number of SG-producing cells was largely abrogated upon introduction of inactivating mutations in N1a and may thus be ascribed to G3BP sequestration. This phenomenon seemed concentration-dependent, noticeable particularly when considering cells with high expression levels of N (**Fig. 5C**), whereas N2b-dependent loss of PKR-induced SGs in wildtype HeLa cells was already observed at very low levels of N (**Fig. 1B**).

**Fig. 5.**
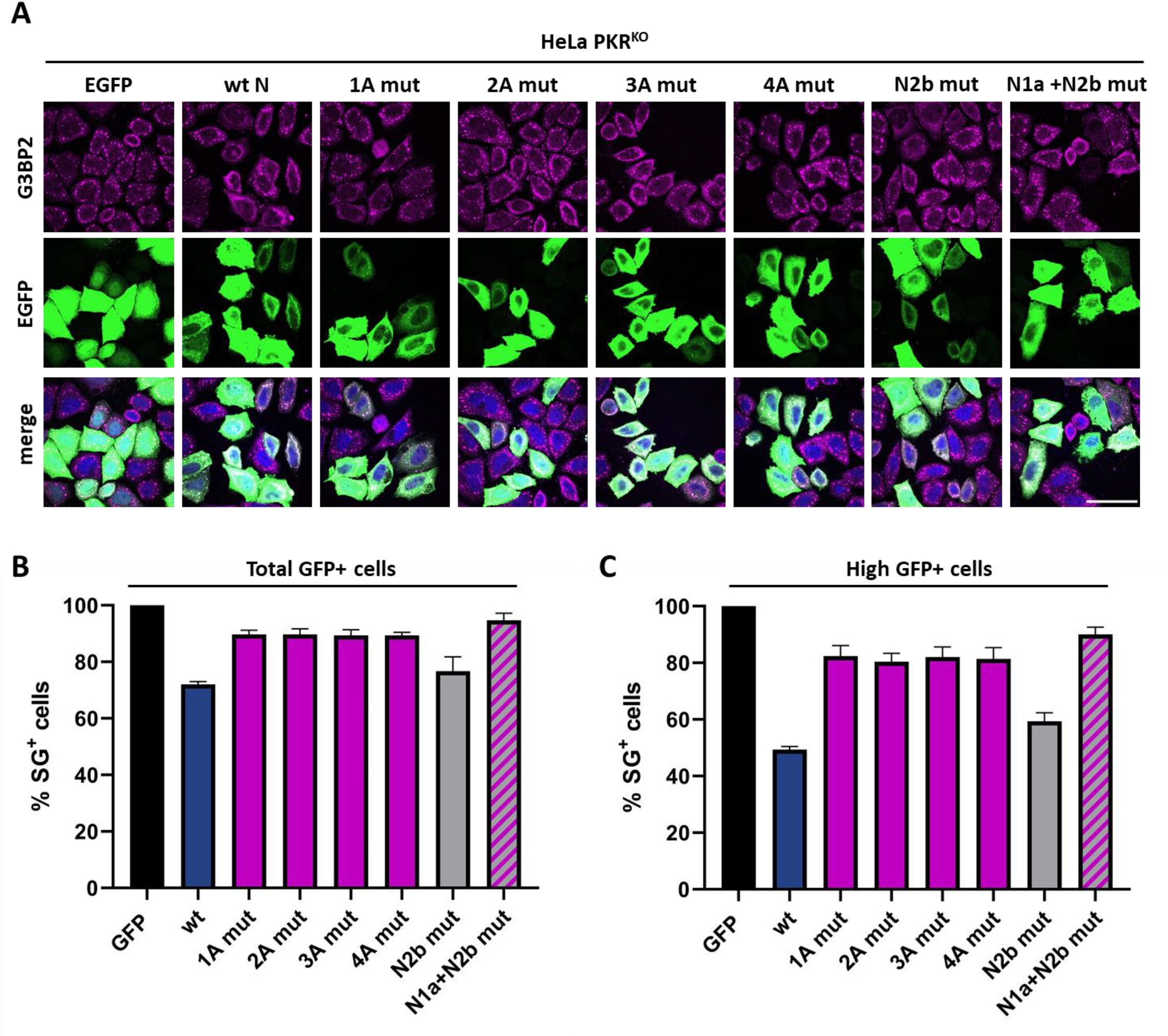
The effect of N-G3BP interaction on arsenite-induced SGs in HeLa PKR^KO^ cells. HeLa PKR^KO^ cells were transfected to express EGFP, SARS-CoV-2 N wt, N mutants defective in G3BP binding (1A-4A mut), N-K^257^A+K^261^A (N2b mut) or a mutant ‘N1a + N2b mut’ with substitutions in both the G3BP binding site (2A mut) and N2b (K^257^A+K^261^A). At 24 hrs post transfection, cells were sodium arsenite-treated to induce HRI-mediated ISR with ensuing formation of SGs. (***A***) Representative (immune)fluorescence images. Scale bar: 50 µm. Quantification of the results based on three independent experiments (***B***) by counting all cells with detectable EGFP fluorescence or (***C***) highly expressing cells exclusively. For a statistical analysis of the results, see **Table S3**.

### Suppression of PKR-induced ISR by coronavirus N proteins is mediated predominantly by N2b

To corroborate our observations, we measured N-mediated rescue of ISR-induced translational arrest also by flow cytometry. To this end, we used a cotransfection assay with red fluorescence protein (RFP) expressed from vector pcDNA-RFP serving as reporter (27). Co-transfection with ISR-inducing plasmid pEGFP-N3 strongly inhibited RFP production (**Fig. 6**, top; see also **Fig. S4**). RFP expression was rescued in cells co-expressing N2b but not by its RNA-binding deficient derivative N2b-K^257^A/K^261^A. RFP expression was also rescued by the intact N proteins of either SARS-CoV-2 or HCoV-229E and to similar levels by SARS-CoV-2 N mutants defective for G3BP1/2 binding (N-R^14^A/I^15^A) or N1b methylation (N-R^95^K) (**Fig. 6**). In contrast, inactivation of N2b through the K^257^A/K^261^A double mutation in the intact N protein of either SARS-CoV-2 or HCoV-229E significantly reduced RFP expression (**Fig. 6**). Intriguingly, however, RFP expression levels were still higher than those observed for EGFP (p-value 0.0014) or N2b-K^257^A/K^261^A (p-value 0.0195). The data suggest that suppression of the PKR-induced ISR by coronavirus N proteins is mediated predominantly by the N2b domain though not exclusively. Apparently, N counteracts translational arrest also through other domains via alternative mechanisms, but this contribution only becomes detectable upon N2b inactivation. Saliently, however, disruption of N-G3BP interactions through substitutions in segment N1a did not have an additive effect when tested in combination with the N2b mutations (**Fig. 6**; panel ‘SARS2 N N1a+N2b mut’).

**Fig. 6.**
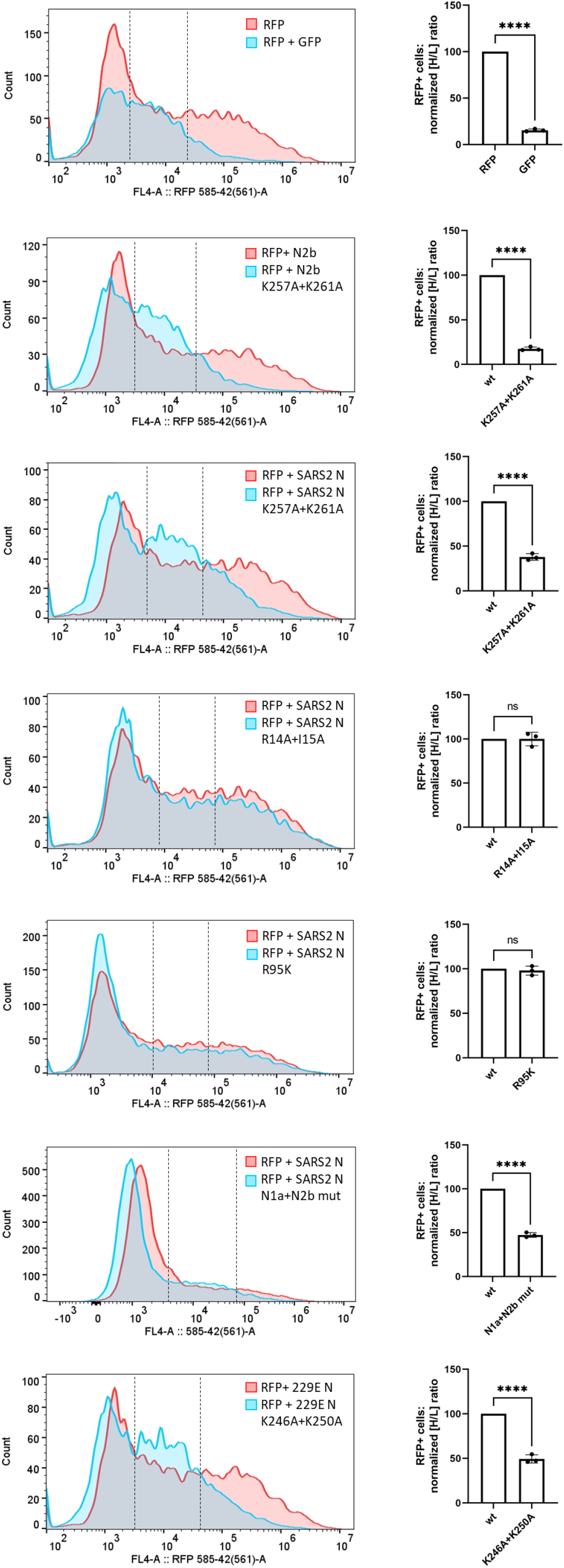
Quantitative assessment of N-mediated rescue of ISR-induced translational arrest. HeLa cells were transfected to express EGFP, SARS-CoV-2 N2b, the full-length N proteins of SARS-CoV-2 and HCoV-229E, and mutants thereof from pEGFP-N3-based expression vectors to induce PKR-activated ISR. The capacity of these proteins to rescue translational repression of red fluorescent protein (RFP) *in trans* or lack thereof was measured by flow cytometry at 24 hr post transfection (left panels) and fluorescence microscopy (see **Fig. S4**). Representative flow cytometry histograms shown. The transfected cells were divided into non-RFP-expressing, low (L) RFP-expressing and high (H) RFP-expressing populations (see dashed lines in histograms). For each mutant protein, the [H/L] ratio was calculated and normalized, with those of the corresponding wildtype proteins set at 100%. The bar graphs show mean values with standard deviations based on three independent experiments (unpaired t-test; ****, P<0.001; ns, non-significant) (right).

### N2b-mediated suppression of the ISR and type I interferon response in virus-infected cells

Whereas the results provide conclusive evidence for ISR suppression upon transient expression of N in transfected cells, the question arises whether this phenomenon also occurs during natural infection and whether it is relevant for evasion of the innate host immune response. Unfortunately, the essential functions of N at multiple stages of the coronavirus life cycle, both during early replication as well was in genome packaging during virion assembly, precludes a straightforward reverse genetics approach. Thus, the construction of recombinant SARS-CoV-2 mutants with an N protein defective in N2b RNA binding was deemed a nonviable option. We therefore resorted to a well-established alternative model based on recombinant encephalomyocarditis virus mutant EMCV-L^zn^, in which the autologous ISR antagonist (the leader protein L) is inactivated. The mutant virus is no longer able to counteract the ISR (63-65) and hence provides a convenient platform to identify and characterize ISR antagonists of other viruses.

Our initial experiments were performed with an EMCV mutant that coded for a chimeric polyprotein with N2b at its N-terminus, downstream of the first five residues of EMCV L^zn^ that are important for efficient IRES-mediated translation (66) and two additional residues encoded by an engineered *Xho*I site. This virus, however, failed to suppress the ISR. We noted that in the fusion protein, two negatively charged residues, E^6^ (from L) and E^8^ (encoded by the *Xho*I sequence) are immediately upstream of N-terminal N2b residues K^248^ and K^249^ (residues 9 and 10 of the chimera) and proximal to critical N2b residues K^257^ and K^261^ **Fig. S5**). Arguing that this might affect N2b RNA binding, we introduced into the EMCV-L^zn^ genome an N-terminally extended N2b with the N2a region serving as a spacer (**Fig. 7A, Fig. S5**). Indeed, in cells, infected with the resulting virus EMCV-L^zn^-N2aN2b^wt^, SG formation was suppressed to 30% of that caused by EMCV-Lzn, i.e. to levels similar to those reported for EMCV derivatives with L replaced by established PKR inhibitor MERS-CoV ns4a (**Fig. 7B, C**). In accordance, in cells infected with the N2aN2b^wt^ virus, levels of phosphorylated PKR were consistently low, comparable to those in wildtype EMCV-infected cells and, as calculated from band densities, 4.5 to 6-fold lower than those in cells infected with EMCV-L^zn-^. Thus, in its activity N2aN2b^wt^ mimicked MERS-CoV ns4a which prevents PKR activation by sequestering dsRNA (27) but differed from AcP10 which inhibits the ISR at a level downstream of PKR (31). Like MERS-CoV ns4a and AcP10, N2aN2b restored replication efficacy of EMCV-L^zn^ to near wildtype levels as based on expression of the viral capsid proteins (**Fig. 7D**).

**Fig. 7.**
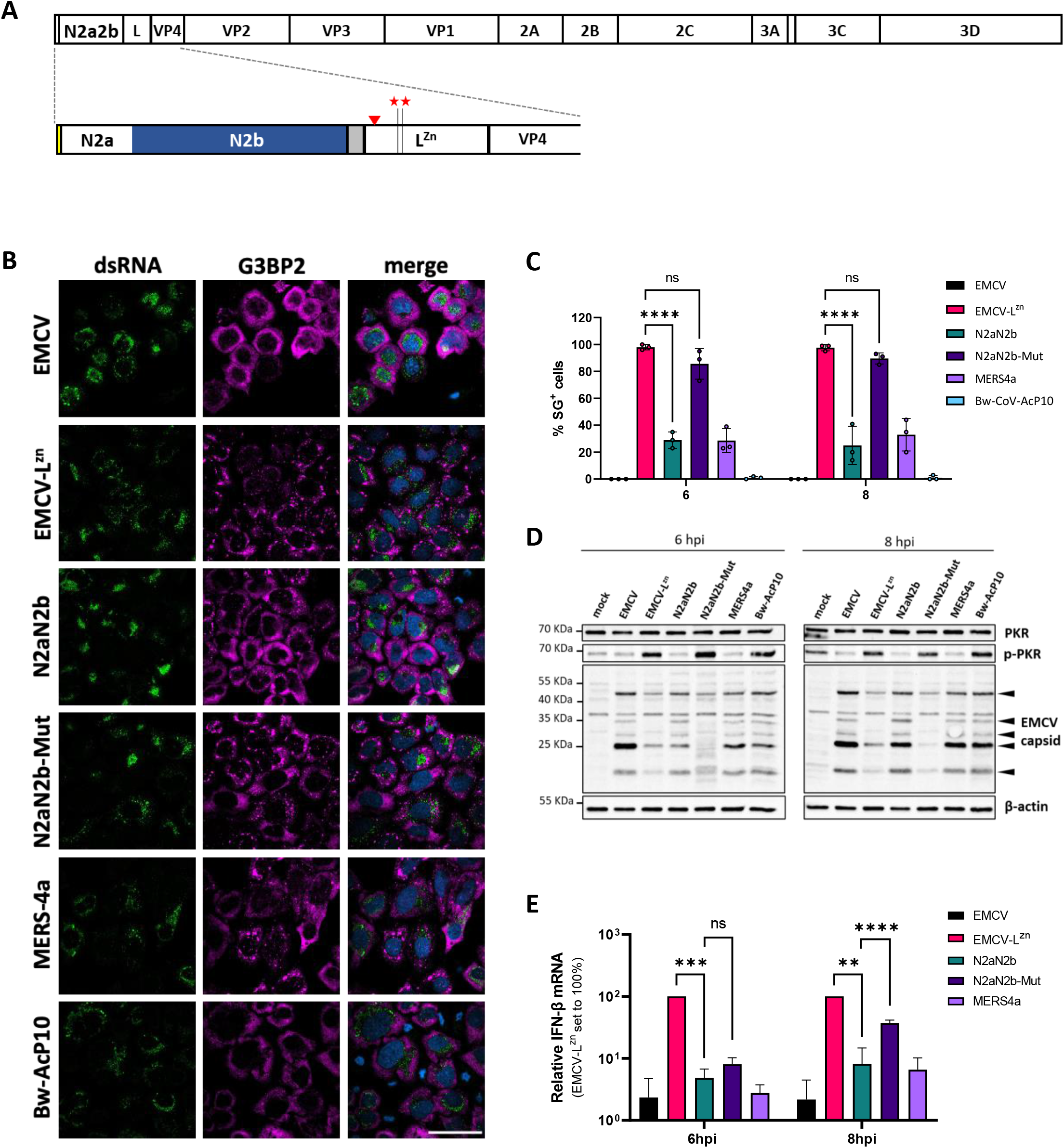
The N2b domain inhibits PKR phosphorylation, prevents SG formation, and reduces the type I interferon response in EMCV infected cells. (***A***) Structure of the polyprotein of recombinant EMCV-L^zn^-N2aN2b. Schematic representation and a close-up of the N-terminal region with mature cleavage products shown as boxes. Red asterisks indicate the locations of mutations (C^19^A/C^22^A) in the Zn-finger domain of leader protein L, which abolish its function as EMCV-autologous antagonist of the host innate immune response. The red arrowhead indicates the location of a newly engineered EMCV 3C cleavage site. The yellow box depicts the native N-terminus of the EMCV polyprotein, i.e. the six N-terminal residues of L and two residues encoded by a synthetic *Xho*I cloning site (see **Fig. S5** for details). The location of the introduced SARS-CoV-2 N2a2b element is given, with the N2b domain in blue and a Strep2 tag in grey. (***B***) SG formation in EMCV-infected cells is prevented by SARS-CoV-2 N2a2b. HeLa cells were infected at MOI 10 with EMCV-L^zn^ recombinant viruses expressing N2a2b or a derivative ‘N2a2b-Mut’ in which the N2b RNA binding site was abolished by Lys^257^Ala and Lys^261^Ala substitutions. Cells infected with parental wildtype EMCV or EMCV-L^zn^ served as controls as were cells infected with recombinant EMCV-L^zn^ derivatives encoding established coronavirus ISR antagonists MERS 4a and BW-Acp10. Cells were fixed at 8 hpi and analyzed by immunofluorescence staining for dsRNA as a marker for infection and for SG scaffold protein G3BP2. (***C***) Percentages of infected cells with stress granules at 6 and 8 hpi. Bar graphs show the means of three independent experiments with more than 200 infected cells counted per sample. Standard deviations indicated by error bars; **** p< 0.0001, ns = not significant (two-way ANOVA with Dunnett post hoc test). (***D***) Western-blot analysis for PKR, phosphorylated PKR (p-PKR), EMCV capsid proteins and β-actin. HeLa cells, mock-infected or infected with recombinant EMCVs at MOI 10, were lysed at 6 or 8 hpi. Images shown are representative of three independent experiments. (***E***) Suppression of type I interferon response in EMCV-infected cells by N2a2b but not N2aN2b-Mut. HeLa cells, mock infected or infected at MOI 10 with the indicated recombinant EMCV-L^zn^ viruses, were lysed at 6 or 8 hpi. Total RNA was isolated and used for RT-qPCR analysis for IFN-β and actin. The IFN-β levels were calculated as fold induction compared to levels in mock-infected cells, after correction for actin mRNA levels, and normalized with EMCV-L^zn^ IFN-β levels set at 100%. Data represent means of three independent experiments. Standard deviations indicated by error bars; statistical significance compared to the results for EMCV-L^zn^ or EMCV-L^zn^-N2aN2b infected cells calculated by two-way ANOVA with the Dunnett post hoc test; ** p<0.01; *** p<0.001; **** p<0.0001.

To test whether N2aN2b, like ns4a, also suppresses the type I interferon response, levels of IFN-β mRNA were determined by qRT-PCR. Indeed, in cells infected with EMCV-L^zn^- N2aN2b^wt^ IFN-β expression was reduced as compared to EMCV-L^zn^ infected cells to levels observed in cells infected with EMCV wt or EMCV-L^zn^-MERS-ns4a (**Fig. 7E**). In contrast, in cells infected with EMCV-L^zn^-N2aN2b-K^257^A+K^261^A, encoding an inactive N2b, there was no detectable suppression of SG formation, PKR was phosphorylated, and viral replication was delayed to similar if not larger extent than in EMCV-L^zn^-infected cells (**Fig. 7B, D**). EMCV-L^zn^- N2aN2b-K^257^A+K^261^A also lost the capacity to suppress the type I IFN response although this was noticeable only at 8 hr post infection (**Fig. 7E**), which may be attributed to the considerable delay in virus replication and a consequential late onset of IFN induction.

We conclude that SARS-CoV-2 N2b functions as an antagonist of the ISR in EMCV-infected cells and can functionally replace the EMCV L protein by acting as a classical PKR antagonist and preventing activation of the ISR by sequestering dsRNA. This mode of action differs from that of beluga whale CoV AcP10 and Aichivirus L, which counteract the ISR by salvaging global translation through competitive inhibition of p-eIF2–eIF2B association (31). The data lend support to the notion that N through N2b may inhibit the ISR and SG formation also in the context of SARS-CoV-2-infected cells.

## DISCUSSION

To successfully establish infection, viruses must subdue the intracellular host defense long enough to complete their replication cycle and vigorously enough to prevent the infected cells from raising the alarm. The detection of dsRNA, an inevitable byproduct of both RNA and DNA virus replication is central to activation of the innate antiviral defense. The activation of RLRs must be averted that would otherwise culminate, through intra- and intercellular signaling, in the production of type I interferons and other proinflammatory cytokines. Also, while host cell protein synthesis should best be inhibited and kept at a minimum, efficient unperturbed production of viral proteins must be ensured by preventing translational suppression due to activation of the OAS/RNaseL pathway and PKR-dependent induction of the ISR. Recent publications implicated SARS-CoV-2 N protein as an inhibitor of SG formation and an RLR antagonist (33-36, 67, 68). Here, we confirmed and extended these observations using a convenient procedure to assess PKR-induced ISR activation based on transfection of ISR-inducing plasmid DNA to drive the expression of putative ISR antagonists genetically fused to EGFP. We conclusively demonstrate that SARS-CoV-2 N is endowed with the ability to inhibit PKR activation, translational arrest and ensuing SG formation. Moreover, we showed that the N protein of human alphacoronavirus 229E (subgenus *Duvinacovirus*) can also prevent translational arrest and suppresses SG formation to a similar extent and via a similar mode of action. These findings suggest that this property may be widely conserved among members of different orthocoronavirus (sub)genera.

To study the mechanism of ISR inhibition, we performed a systematic deletion and site-directed mutational analysis of SARS-CoV-N. We found that inhibition of SG formation was lost upon deletion of N2b in accordance with findings of Zheng et *al*., (35). However, we crucially extend their observations by showing that N2b when expressed in isolation, is sufficient to impair transfection-induced ISR-mediated translational shut off as well as the formation of SGs. Apparently, N2b counteracts the ISR by binding dsRNA. Ala substitution of conserved Lys residues (Lys^257^ and/or Lys^261^) in the putative RNA-binding cleft of N2b that abolished dsRNA binding also abolished ISR inhibition. Importantly, these substitutions also caused the intact N proteins of SARS-CoV-2 and HCoV-229E to lose their activity as ISR antagonists. The observations thus strongly support a model in which coronavirus N proteins through their N2b domain prevent activation of the ISR by PKR by sequestering dsRNA, like the other viral PKR inhibitors IAV ns1 and MERS-CoV ns4a that were used as controls throughout.

The N protein plays a crucial role during the very early steps of the CoV infection cycle (55, 69-71) and is indispensable for virus morphogenesis. Evidently, this impedes a direct mutational reverse genetics approach to study the occurrence and relevance of N-mediated inhibition of the ISR in CoV-infected cells. However, by using a mutant EMCV as a platform, we showed that an extended protein N2aN2b can functionally replace the autologous enterovirus L protein and can salvage the EMCV replication defect caused by L inactivation, like we previously showed for MERS-CoV ns4a and BWCoV AcP10 (27, 31). Like MERS-CoV ns4a, but unlike BWCoV AcP10 -which blocks the ISR downstream of PKR-, N2aN2b prevented PKR phosphorylation apparently via N2b-mediated RNA binding. The findings provide proof of principle that N2b can promote viral replication by preventing PKR and ensuing ISR activation, supporting the notion that N through its N2b domain may do so as well during CoV infection.

Adding relevance to our observations, selection for enhanced innate immune escape apparently favored SARS-CoV-2 variants with increased N expression (72). Moreover, a three-nucleotide change in Gamma also present in Alpha and Omicron variants of concern created a new (cryptic) transcription regulating sequence to drive the synthesis of a novel subgenomic mRNA species from which a truncated N-protein, N*, may be translated corresponding with the C-terminus of the protein and initiating at in frame Met^210^ (72, 73). If N* is indeed expressed, our findings strongly support a role in dsRNA sequestration and innate antagonism.

The native coronavirus N proteins are subject to extensive posttranslational modifications in domains and segments other than N2b. Some have been implicated in liquid-liquid phase separation (62, 74), RNA binding (33), SG formation and inhibition of innate immunity (33, 62), prompting the question whether these modifications promote or hinder its function as an ISR antagonist. Reportedly, methylation of Arg^95^ in the SARS-CoV-2 N1b segment by host protein arginine methyltransferases is required for RNA binding and for the inhibition of arsenite-induced formation of G3BP1-containing SGs (33). In our hands, however, an N mutant with Arg^95^Lys substitution, described to block N-mediated suppression of SGs (33), still prevented PKR-induced ISR-associated translational repression to identical extent as the wildtype protein and likewise prevented SG formation. In another recent study, Wang et *al*. (62) reported suppression of MAVS signaling by intact SARS-CoV-2 N protein but not by an N derivative from which N2b had been deleted. They attributed this to N-induced and N2b-dependent liquid-liquid phase separation (LLPS) subject to acetylation of N3 residue Lys^375^. In accordance, we find that in EMCV-infected cells, N2aN2b, like MERS4a, suppresses β-interferon (IFN-β) expression and show that this activity is blocked by mutations that abolish N2b-mediated RNA binding. In apparent contrast to the observations of Wang et *al*. (62), however, we find that acetylation of Lys^375^ is not essential for RNA binding by N2b and that Lys^375^Arg substitution does not detectably affect inhibition of the PKR-induced ISR by the intact N protein. Our findings suggest that N, through the autonomous activity of the N2b domain, acts not only as a classical PKR inhibitor to thwart ISR activation but may also prevent RLR activation and thus block induction of the type I IFN response, whether directly by shielding dsRNA or indirectly by preventing SGs to act as platform for immune signaling.

The interaction between N and SG scaffold proteins G3BP1 and -2 mediated through the N1a ФxFG motif (33, 37-40), was recently concluded to be the main determinant in SG disassembly (38). Our current findings would seem to be at odds with this view, but in accordance with observations by these authors, we did observe that functional disruption of the critical ФxFG motif abrogated inhibition of SG formation in arsenite-treated HeLa PKR^KO^ cells. Our findings, however, would suggest that sequestration of the G3BPs to prevent assembly of SG or to promote their disassembly requires high expression levels of N. Importantly, N proteins with intact N1a domain but with mutations in N2b that abolish RNA binding failed to prevent SG formation in HeLa wt cells. Also, the 229E N protein lacks a ФxFG motif, yet inhibits formation of SGs via the same N2b-dependent mechanism as SARS-CoV-2 N. The findings may be reconciled, however, by assuming that sarbecovirus N proteins have been evolutionary selected to interfere with G3BP function, possibly even beyond the formation and function of SGs (75), through distinct mechanisms at multiple levels during different stages of the viral life cycle, mediated by different domains and regulated by distinct posttranslational modifications, and perhaps subject to protein distribution and availability of viral genomes for encapsidation. Such subtleties may be missed in over-expression experiments and overshadowed by the robust inhibition of the PKR-triggered ISR by N2b. Hence, in coronavirus-infected cells, N1a-mediated G3BP-N association (38, 40, 54) and N2b-mediated dsRNA binding (36) may well act in concert to hamper SG assembly and to impede recruitment and SG-facilitated activation of RLRs and PKR (20-23). Indeed, this would be consistent with observations by others (35, 36) of SARS-CoV-2 N suppressing SG-associated RLR activation and inhibiting induction of type I IFNs by targeting G3BP1. The relative importance of dsRNA shielding, G3BP recruitment and other activities of N for suppression of innate immunity during coronavirus infection clearly deserves further study.

## MATERIALS AND METHODS

### Cell lines

HeLa-R19, HeLa R19 PKR^KO^ (27), A549 and BHK21 cells were maintained in Dulbecco’s Modified Eagle’s Medium (DMEM) supplemented with 10% (V/V) fetal calf serum (FCS) and 100 units/ml penicillin and streptomycin. All cell lines were routinely tested negative for mycoplasma contamination.

### Recombinant EMCV viruses

Recombinant EMCV viruses were derived from the pM16.1-derived pStrep2-VFETQG-Zn-M16.1 infectious clone (27, 76). This vector carries a Zn-finger mutation to inactivate EMCV L as well as coding sequences for a Strep2-tag and a synthetic optimized 3C^pro^ recognition site (VFETQG). The coding sequences for N2b and N2a2b were PCR amplified from SARS-CoV-2-derived cDNA (sequences deposited in EVA database; Ref-SKU: 026V-03883) and inserted into XhoI/NotI-digested pStrep2-VFETQG-Zn-M16.1 yielding pStrep2-SARS-CoV-2-N2b-VFETQG-Zn-M16.1 and pStrep2-SARS-CoV-2-N2aN2b-VFETQG-Zn-M16.1, respectively. Mutations to substitute N2b Lys^257^ and Lys^261^ by Ala were introduced by Q5 side-directed mutagenesis (New England Biolabs, NEB) generating pStrep2-SARS-CoV-2-N2aN2b K^257^A/K^261^A-VFETQG-Zn-M16.1. The vectors were linearized with BamHI, used for *in vitro* transcription with the RiboMAX kit (Promega) and the resulting RNA was purified using the NucleoSpin RNA mini kit (Machery-Nagel) following the manufacturer’s protocol. Viruses were recovered by transfecting BHK-21 cells, grown to subconfluency in T125 flasks, with 1.5 µg of the run-off RNA transcripts using Lipofectamine2000 (Invitrogen) transfection reagent according to the manufacturer’s protocol. After 2-4 days, when total cytopathic effect was apparent, the cultures were subjected to three freeze-thaw cycles, cell debris was pelleted at 4,000x*g* for 15 minutes and virus was concentrated from the supernatants by ultracentrifugation though a 30% sucrose cushion at 140,000x*g* for 16 hrs at 4°C in a SW32Ti rotor. Virus pellets were resuspended in phosphate-buffer saline (PBS). The viruses were characterized by isolating viral RNA from 150-µL aliquots of the cell culture supernatant with the NucleoSpin RNA Virus kit (Macherey-Nagel) followed by conventional RT-PCR and bidirectional Sanger sequence analysis of the inserted SARS-CoV-2 sequences and flanking regions. Viral titers, determined by endpoint titration and calculated by the Spearman-Kaerber formula, are averages from three independent experiments.

### Plasmids for eukaryotic and prokaryotic expression

Eukaryotic expression plasmids were constructed by cloning PCR-amplified sequences of interests, flanked by NheI and BamHI restriction sites, into NheI/BamHI digested vector pEGFP-N3 (ClonTech) such that the encoded viral proteins are C-terminally fused to enhanced green fluorescent protein (EGFP). pcDNA-RFP was purchased from Addgene.

Procaryotic expression plasmid pGEX2T-Hisx6-N2b encoding N-terminally His-tagged SARS-CoV-2 N2b was created by linearizing vector pGEX2T, amplifying the N2b coding sequences by Q5-PCR, and assembling the fragments by NEBuilder HiFi DNA Assembly (NEB) according to the manufacturer’s instructions. Mutations (N2b K^257^A, N2b K^261^A and N2b K^257^A+K^261^A) were generated by Q5 site-directed mutagenesis (New England Biolabs, NEB). All constructs were sequenced to confirm their integrity. There were no major differences in transfection efficiency (**Fig. S6**)

### Immunofluorescence assay

HeLa-R19 wt, HeLa-R19 PKR^KO^ cells and A549 cells were seeded onto 12 mm glass cover slips in 24-well plates (Corning Costar) at 5×10^4^ cells and grown for 24 hr. They were then transfected with 500 ng total DNA/well using Lipofectamine2000 (Invitrogen) according to the instructions of the manufacturer. At 24 hr post transfection, cells were either left untreated or treated with 500 µM sodium arsenite (NaAsO_2_, Riedel-de Haën) for 30 min at 37°C and subsequently fixed in PBS + 3.7% paraformaldehyde (PFA).

Vero E6-TMPRSS2 cells were infected with SARS-CoV-2 Wuhan (D614G) or Omicron BA.1 variants at a multiplicity of infection (MOI) of 5 TCID_50_/cell and HeLa-R19 cells were infected with recombinant EMCV viruses at an MOI of 10 TCID_50_/cell and incubated for times indicated in the text. Cells were fixed with paraformaldehyde (3.7% in PBS), incubated with PBS + 0.1% glycine for 10 min to quench the fixative, permeabilized with 0.1% Triton X-100/PBS+s for 10 min at RT and subsequently blocked in blocking buffer (PBS + 1% BSA + 0.1% Tween-20) for 30min in a dark humidified chamber at 37°C. The cells were then incubated in blocking buffer containing mouse anti-dsRNA (1:1000; English & Scientific Consulting), goat anti-eIF3η (1:200, SantaCruz) and rabbit anti-G3BP2 (1:200; Bethyl Laboratories) for 1h at room temperature (RT). After three washing steps with PBS+0.1%Tween-20, cells were incubated with secondary antibody donkey anti-mouse Cy2 (1:100; The Jackson Laboratory), donkey anti-goat Alexa594 (1:200; Invitrogen), donkey anti-rabbit Alexa647 (1:200; Invitrogen) in block buffer for 1h at RT. The cells were then washed three times with PBS+0.1%Tween-20, once with distilled water and mounted on glass microscope slides in ProLong Diamond Antifade (Invitrogen) mounting medium. Cells were examined by conventional wide-field (Olympus) and confocal immunofluorescence microscopy (Nikon A1R) in most cases also in a blinded fashion by a second independent observer.

### Flowcytometry analysis

HeLa cells were seeded in a 12-well cluster (10^5^ cells/well) and, after overnight incubation, transfected with the indicated plasmids (500 ng well; 250 ng/plasmid) using Lipofecatmine2000 (Invitrogen) according to the manufacturer’s instructions. At 24 hr post transfection, cells were detached with trypsin, washed once PBS, and fixed in PBS, 3.7% PFA for 10 min. The samples were washed once in FACS buffer (PBS + 0.02% Na-azide and 0.5% BSA) and kept in FACS buffer at 4°C. RFP fluorescence intensity was recorded with a CytoFLEX LX flow cytometer (Beckman Coulter) and the data were analyzed by FlowJo™ v10 software (BD Biosciences). Samples were gated for live single cell populations and then gated for negative, low (L) and high (H) RFP expressing cells. For bar graphs [H/L] ratios were normalized with the ratio calculated for the relevant wildtype protein set at 100%.

### Electrophoretic Mobility Shift Assay (EMSA)

To purify the wildtype N2b domain and its mutated derivatives, *E. coli* BL21 cells (Sigma-Aldrich), transformed with pGEX2T-Hisx6-N2b or its mutated derivatives, were grown in LB medium containing ampicillin (50 μg/ml) at 37°C until the optical density at 600 nm (OD_600nm_) reached 0.3. The temperature was then reduced to 18°C, and when the OD_600nm_ reached 0.5, protein expression was induced with 0.5 mM IPTG. Following expression for 16 hrs, the cells were harvested by centrifugation and resuspended in lysis buffer (100 mM Tris-HCl pH=8, 300 mM NaCl, 10 mM imidazole, 0.1% Triton X-100, 5% Glycerol) complemented with lysozyme (0.25 mg/ml; Merk) and cOmplete Protease Inhibitor (Roche). The samples were sonicated and centrifuged at 15,000*g* for 45 min at 4°C to pellet cell debris. The cleared lysates were passed through a 0.45-μm filter and incubated with 1 ml of Ni-NTA (nitrilotriacetic acid) resin (Thermo Scientific) at 4°C overnight (O/N) on a roller. The beads were washed twice with wash buffer (100mM Tris-HCl (pH 8.0), 300mM NaCl, 10mM Imidazole) and then eluted with a buffer consisting of 100 mM Tris-HCl, 300 mM NaCl, and 500 mM imidazole (pH 8). The eluted proteins were subjected to dialysis in 20mM Tris-HCl, 150mM NaCl and flash-frozen in 20-μl aliquots.

For the EMSA, 1 µM of a 32-mer single stranded RNA oligonucleotides corresponding to SARS-CoV-2 (GenBank accession no. MZ558051.1) sequence 5′-CGAGGCCACGCGGAGUACGAUCGAGGGUACAG-3′ and a scrambled version thereof, 5’-GGCACGGAGUAUACCGGACGAGCGGAACGGCU-3’, each were mixed 1 :1 with their respective complimentary RNA oligonucleotides in denaturing buffer (10 mM NaH_2_PO_4_·H_2_O, 50 mM NaCl, 1 mM EDTA, 0.01% NaN_3,_ pH 7.4 at 25°C), incubated at 90°C for 4 min and then allowed to anneal at RT for 30 min. For the ssRNA sample preparation, the oligonucleotide was diluted in denaturing buffer, incubated at 90°C for 30 sec to destroy possible secondary structures and rapidly cooled on ice. Proteins were diluted in protein buffer (20mM Tris-HCl (pH=8), 150mM NaCl) to a concentration of 100 ng/µl and mixed in 10X, 20X or 40X fold molar excess with 10 ng ssRNA or 10 ng dsRNA in binding buffer (20 mM Tris-HCl, 100 mM NaCl, 1 mM EDTA, 1mM TCEP, 0.02% Tween-20, pH 7.0 at 25°C) in 5 μl reaction volumes. The samples were incubated for 30 min on ice, then supplemented with glycerol to a final concentration of 5% (v/v) and separated in ultrathin (10 ml; 75 × 50 mm) RNase-free 1% agarose gels in 0.5x TB buffer (45 mM Tris base, 45 mM boric acid, pH 8.2-8.5) at 200V. RNAs were stained by incubating the gels in 50 ml 2X Invitrogen™ SYBR™ Gold nucleic acid gel stain (ThermoFisher) in 0.5x TB buffer, diluted from a 10.000X concentrate, for 15 min under agitation and de-stained for 21 min with 0.5 X TB buffer refreshing the buffer 3 times. Stained RNA was visualized with the Gel Doc System (BioRad).

### Western-blot analysis

HeLa R19 wt and PKR^KO^ cells were seeded in 6-well clusters (4×10^5^ cells/well) and after a 16 hr recovery were infected with recombinant EMCVs at MOI 10 or tranfected with plasmids as indicated in the text. The infection was allowed to proceed for 6 or 8 hrs and transfections for 24 hrs. Cells were then released by trypsin and lysed in ice-cold lysis buffer (Tris-HCl pH 8.0, 50mM, NaCl 150mM, EDTA 1mM, NP40 1%, cOmplete Mini Protease Inhibitor Cocktail (Roche) and phosphatase inhibitor PhosSTOP (Roche)) for 30 min at 4°C under constant agitation. Cell debris was pelleted for 20 min 12000 rpm at 4°C. Cleared lysates were harvested and protein concentrations determined by the Pierce BCA assay (ThermoFisher Scientific) according to the manufacturer’s instructions. Samples of each lysate, corresponding to 100 µg protein, were separated by SDS-PAGE in reducing 10% polyacrylamide gels. The proteins were then blotted onto 0.2 µm nitrocellulose membranes by wet electrophoretic transfer or semi-dry transfer. Membranes were washed three times in TBST (20 mM Tris, 150 mM NaCl + 0.1% Tween-20), 5 min each, and incubated in blocking buffer (TBST + 2% BSA) for 1h at RT. Membranes were successively incubated with primary antibodies diluted in blocking buffer (mouse anti-G3BP1, 1:4000, BD Biosciences, rabbit anti-GFP, 1:1000, ThermoFisher, mouse anti-PKR, 1:1000, BD Biosciences; rabbit anti-PKR-P, 1:1000, Abcam; mouse anti-βactin, 1:5000, Invitrogen; rabbit anti-mengovirus capsid, 1:1000, kindly provided by Prof. Ann Palmenberg) for 16 hr at 4°C, and then for 30 min at RT with goat-α-mouse-IRDye680 (Li-COR, 1:15000) or goat-α-rabbit-IRDye800 (Li-COR, 1:15000) diluted in blocking buffer. Between and after the incubations, the membranes were washed, thrice each time, with TBST. Finally, membranes were washed once with PBS and scanned using an Odyssey Imager (Li-COR). ImageJ was used to quantitate and compare density of bands after correction with beta-actin as loading control.

### Co-immunoprecipitation (Co-IP) assay

HeLa R19 wt and PKR^KO^ cells were seeded in 6-well clusters (4×10^5^ cells/well) and, after a 16 hr recovery, transfected with the indicated plasmids. At 24 hrs post transfection, cells were washed once in PBS, released by trypsin and incubated in ice-cold lysis buffer (Tris-HCl pH 8.0, 50mM, NaCl 150mM, EDTA 1mM, NP40 1%, cOmplete Mini Protease Inhibitor Cocktail (Roche) and phosphatase inhibitor PhosSTOP (Roche)) for 30 min on ice. The cell lysates were cleared in a microcentrifuge for 10 min at 12000 rpm at 4°C. Supernatants were harvested and incubated with 25 µl of preequilibrated GFP-Trap agarose beads (ChromoTek) for 1h at 4°C with end-over-end rotation. Beads were then washed 3 times with wash buffer (Tris-HCl pH 8.0, 50mM, NaCl 150mM, EDTA 1mM). Bound proteins were eluted in 80 µl 2x SDS-sample buffer, separated by SDS-PAGE in reducing 8% polyacrylamide gels and analyzed by Western blotting.

### RT-qPCR analysis

HeLa R19 cells, seeded in 24-well clusters (5×10^4^ cells/well), were inoculated with recombinant EMCVs at MOI 10 as above. At 6 or 8 hrs post infection cells were lysed and cellular RNA was isolated using the total RNA isolation kit (Machery-Nagel) according to manufacturer’s instructions. Reverse transcription was set up using TaqMan Reverse Transcription Reagents (Applied Biosystem) as indicated in the manufacturer’s instructions. qPCR analysis of human IFN-β, human actin mRNA and EMCV viral RNA was performed mixing the Fast SYBR green Master Mix (ThermoFisher) with cDNA and 1µM of the forward and reverse corresponding primers: IFN-β (5’-ATGACCAACAAGTGTCTCCTCC-3’ and 5’-GCTCATGGAAAGAGCTGTAGTG-3’), actin (5’-CCTTCCTGGGCATGGAGTCCTG-3’ and 5’GGAGCAATGATCTTGATCTTC-3’) and EMCV RNA (5’-TCTGTTCTGCCTGTGTTTG-3’ and 5’-AAAGAAGAGGGTGCCGAAAT-3’). Amplification occurred in a Roche Light Cycler using the following program: polymerase activation (95°C for 5min), amplification (45 cycles: 95°C for 10 sec, 60°C for 5 sec, 72°C for 30 sec), melting curve (95°C for 5 sec, 65°C for 1 min) and cooling (40°C for 10 sec). The experiments were carried out in triplex for each data point. The relative quantification of the IFN-β gene expression was determined using the 2^-ΔΔCt^ method (77), then the relative IFN-β mRNA levels were normalized to the EMCV-zn IFN-β mRNA level set as 100.

## ACKNOWLEDGMENTS AND FUNDING SOURCES

We would like to thank Huib H. Rabouw for helpful comments and discussions, Richard Wubbolts and Ester van t Veld from the Center for Cell Imaging for the microscopy support and Ger Arkesteijn and Estefania Lozano Andres from the Flow Cytometry and Cell Sorting Facility for the flow cytometry support.

This work was supported by the European Union (Horizon 2020 Marie Skłodowska-Curie ETN “INITIATE”, grant agreement number 813343) and by the Dutch Research Council NWO OCENW.KLEIN.344.

## REFERENCES

1. S. Hur, Double-Stranded RNA Sensors and Modulators in Innate Immunity. Annual Review of Immunology 37, 349–375 (2019).

2. D. H. Wreschner, J. W. McCauley, J. J. Skehel, I. M. Kerr, Interferon action—sequence specificity of the ppp(A2′p)nA-dependent ribonuclease. Nature 289, 414–417 (1981).

3. J. B. Andersen, K. Mazan-Mamczarz, M. Zhan, M. Gorospe, B. A. Hassel, Ribosomal protein mRNAs are primary targets of regulation in RNase-L-induced senescence. RNA Biol 6, 305–315 (2009).

4. Y. Li et al., Activation of RNase L is dependent on OAS3 expression during infection with diverse human viruses. Proceedings of the National Academy of Sciences 113, 2241–2246 (2016).

5. J. Donovan, S. Rath, D. Kolet-Mandrikov, A. Korennykh, Rapid RNase L-driven arrest of protein synthesis in the dsRNA response without degradation of translation machinery. Rna 23, 1660–1671 (2017).

6. J. M. Burke, S. L. Moon, T. Matheny, R. Parker, RNase L Reprograms Translation by Widespread mRNA Turnover Escaped by Antiviral mRNAs. Mol Cell 75, 1203-1217.e1205 (2019).

7. S. L. Schwartz, G. L. Conn, RNA regulation of the antiviral protein 2’-5’-oligoadenylate synthetase. Wiley Interdiscip Rev RNA 10, e1534 (2019).

8. A. G. Hinnebusch, The scanning mechanism of eukaryotic translation initiation. Annu Rev Biochem 83, 779–812 (2014).

9. J. L. Llácer et al., Conformational Differences between Open and Closed States of the Eukaryotic Translation Initiation Complex. Mol Cell 59, 399–412 (2015).

10. M. Yoneyama, M. Jogi, K. Onomoto, Regulation of antiviral innate immune signaling by stress-induced RNA granules. The Journal of Biochemistry 159, 279–286 (2016).

11. Y. Wu, Z. Zhang, Y. Li, Y. Li, The Regulation of Integrated Stress Response Signaling Pathway on Viral Infection and Viral Antagonism. Frontiers in Microbiology 12 (2022).

12. N. Kedersha et al., Evidence that ternary complex (eIF2-GTP-tRNA(i)(Met))-deficient preinitiation complexes are core constituents of mammalian stress granules. Mol Biol Cell 13, 195–210 (2002).

13. M. D. Panas, P. Ivanov, P. Anderson, Mechanistic insights into mammalian stress granule dynamics. J Cell Biol 215, 313–323 (2016).

14. S. Hofmann, N. Kedersha, P. Anderson, P. Ivanov, Molecular mechanisms of stress granule assembly and disassembly. Biochim Biophys Acta Mol Cell Res 1868, 118876 (2021).

15. H. Tourrière et al., The RasGAP-associated endoribonuclease G3BP assembles stress granules. J Cell Biol 160, 823–831 (2003).

16. N. Gilks et al., Stress granule assembly is mediated by prion-like aggregation of TIA-1. Mol Biol Cell 15, 5383–5398 (2004).

17. J. R. Wheeler, T. Matheny, S. Jain, R. Abrisch, R. Parker, Distinct stages in stress granule assembly and disassembly. eLife 5, e18413 (2016).

18. J. Guillén-Boixet et al., RNA-Induced Conformational Switching and Clustering of G3BP Drive Stress Granule Assembly by Condensation. Cell 181, 346-361.e317 (2020).

19. P. Yang et al., G3BP1 Is a Tunable Switch that Triggers Phase Separation to Assemble Stress Granules. Cell 181, 325-345.e328 (2020).

20. K. Onomoto et al., Critical role of an antiviral stress granule containing RIG-I and PKR in viral detection and innate immunity. PLoS One 7, e43031 (2012).

21. J. S. Yoo et al., DHX36 enhances RIG-I signaling by facilitating PKR-mediated antiviral stress granule formation. PLoS Pathog 10, e1004012 (2014).

22. L. C. Reineke, R. E. Lloyd, The stress granule protein G3BP1 recruits protein kinase R to promote multiple innate immune antiviral responses. J Virol 89, 2575–2589 (2015).

23. W. Yang et al., G3BP1 inhibits RNA virus replication by positively regulating RIG-I-mediated cellular antiviral response. Cell Death & Disease 10, 946 (2019).

24. X. Deng et al., Coronavirus nonstructural protein 15 mediates evasion of dsRNA sensors and limits apoptosis in macrophages. Proc Natl Acad Sci U S A 114, E4251–e4260 (2017).

25. E. Kindler et al., Early endonuclease-mediated evasion of RNA sensing ensures efficient coronavirus replication. PLoS Pathog 13, e1006195 (2017).

26. J. Zhao et al., Coronavirus Endoribonuclease Ensures Efficient Viral Replication and Prevents Protein Kinase R Activation. J Virol 95 (2020).

27. H. H. Rabouw et al., Middle East Respiratory Coronavirus Accessory Protein 4a Inhibits PKR-Mediated Antiviral Stress Responses. PLoS Pathog 12, e1005982 (2016).

28. K. Nakagawa, K. Narayanan, M. Wada, S. Makino, Inhibition of Stress Granule Formation by Middle East Respiratory Syndrome Coronavirus 4a Accessory Protein Facilitates Viral Translation, Leading to Efficient Virus Replication. J Virol 92 (2018).

29. J. M. Thornbrough et al., Middle East Respiratory Syndrome Coronavirus NS4b Protein Inhibits Host RNase L Activation. mBio 7, e00258 (2016).

30. S. A. Goldstein et al., Lineage A Betacoronavirus NS2 Proteins and the Homologous Torovirus Berne pp1a Carboxy-Terminal Domain Are Phosphodiesterases That Antagonize Activation of RNase L. J Virol 91 (2017).

31. H. H. Rabouw et al., Inhibition of the integrated stress response by viral proteins that block p-eIF2–eIF2B association. Nature Microbiology 5, 1361–1373 (2020).

32. B. Gao et al., Inhibition of anti-viral stress granule formation by coronavirus endoribonuclease nsp15 ensures efficient virus replication. PLoS Pathog 17, e1008690 (2021).

33. T. Cai, Z. Yu, Z. Wang, C. Liang, S. Richard, Arginine methylation of SARS-Cov-2 nucleocapsid protein regulates RNA binding, its ability to suppress stress granule formation, and viral replication. J Biol Chem 297, 100821 (2021).

34. L. Luo et al., SARS-CoV-2 nucleocapsid protein phase separates with G3BPs to disassemble stress granules and facilitate viral production. Sci Bull (Beijing) 66, 1194–1204 (2021).

35. Z.-Q. Zheng, S.-Y. Wang, Z.-S. Xu, Y.-Z. Fu, Y.-Y. Wang, SARS-CoV-2 nucleocapsid protein impairs stress granule formation to promote viral replication. Cell Discovery 7, 38 (2021).

36. H. Liu et al., SARS-CoV-2 N Protein Antagonizes Stress Granule Assembly and IFN Production by Interacting with G3BPs to Facilitate Viral Replication. J Virol 96, e0041222 (2022).

37. D. E. Gordon et al., A SARS-CoV-2 protein interaction map reveals targets for drug repurposing. Nature 583, 459–468 (2020).

38. T. Kruse et al., Large scale discovery of coronavirus-host factor protein interaction motifs reveals SARS-CoV-2 specific mechanisms and vulnerabilities. Nature Communications 12, 6761 (2021).

39. J. Li et al., Virus-Host Interactome and Proteomic Survey Reveal Potential Virulence Factors Influencing SARS-CoV-2 Pathogenesis. Med (N Y) 2, 99-112.e117 (2021).

40. M. Biswal, J. Lu, J. Song, SARS-CoV-2 Nucleocapsid Protein Targets a Conserved Surface Groove of the NTF2-like Domain of G3BP1. J Mol Biol 434, 167516 (2022).

41. K. H. Mellits, T. Pe’ery, L. Manche, H. D. Robertson, M. B. Mathews, Removal of double-stranded contaminants from RNA transcripts: synthesis of adenovirus VA RNA 1 from a T7 vector. Nucleic Acids Research 18, 5401–5406 (1990).

42. F. Terenzi et al., The antiviral enzymes PKR and RNase L suppress gene expression from viral and non-viral based vectors. Nucleic Acids Research 27, 4369–4375 (1999).

43. T. Gantke et al., Ebola virus VP35 induces high-level production of recombinant TPL-2-ABIN-2-NF-κB1 p105 complex in co-transfected HEK-293 cells. Biochem J 452, 359–365 (2013).

44. K. R. Hurst, C. A. Koetzner, P. S. Masters, Identification of in vivo-interacting domains of the murine coronavirus nucleocapsid protein. J Virol 83, 7221–7234 (2009).

45. K. R. Hurst, R. Ye, S. J. Goebel, P. Jayaraman, P. S. Masters, An interaction between the nucleocapsid protein and a component of the replicase-transcriptase complex is crucial for the infectivity of coronavirus genomic RNA. J Virol 84, 10276–10288 (2010).

46. C. K. Chang, M. H. Hou, C. F. Chang, C. D. Hsiao, T. H. Huang, The SARS coronavirus nucleocapsid protein--forms and functions. Antiviral Res 103, 39–50 (2014).

47. Q. Ye, A. M. V. West, S. Silletti, K. D. Corbett, Architecture and self-assembly of the SARS-CoV-2 nucleocapsid protein. Protein Sci 29, 1890–1901 (2020).

48. T. Y. Peng, K. R. Lee, W. Y. Tarn, Phosphorylation of the arginine/serine dipeptide-rich motif of the severe acute respiratory syndrome coronavirus nucleocapsid protein modulates its multimerization, translation inhibitory activity and cellular localization. Febs j 275, 4152–4163 (2008).

49. E. Nikolakaki, T. Giannakouros, SR/RS Motifs as Critical Determinants of Coronavirus Life Cycle. Front Mol Biosci 7, 219 (2020).

50. C. A. Koetzner, K. R. Hurst-Hess, L. Kuo, P. S. Masters, Analysis of a crucial interaction between the coronavirus nucleocapsid protein and the major membrane-bound subunit of the viral replicase-transcriptase complex. Virology 567, 1–14 (2022).

51. R. Molenkamp, W. J. Spaan, Identification of a specific interaction between the coronavirus mouse hepatitis virus A59 nucleocapsid protein and packaging signal. Virology 239, 78–86 (1997).

52. L. Kuo, C. A. Koetzner, K. R. Hurst, P. S. Masters, Recognition of the murine coronavirus genomic RNA packaging signal depends on the second RNA-binding domain of the nucleocapsid protein. J Virol 88, 4451–4465 (2014).

53. K. R. Hurst, C. A. Koetzner, P. S. Masters, Characterization of a critical interaction between the coronavirus nucleocapsid protein and nonstructural protein 3 of the viral replicase-transcriptase complex. J Virol 87, 9159–9172 (2013).

54. W. Huang et al., Molecular determinants for regulation of G3BP1/2 phase separation by the SARS-CoV-2 nucleocapsid protein. Cell Discovery 7, 69 (2021).

55. C. Y. Chen et al., Structure of the SARS coronavirus nucleocapsid protein RNA-binding dimerization domain suggests a mechanism for helical packaging of viral RNA. J Mol Biol 368, 1075–1086 (2007).

56. M. Yang et al., Structural Insight Into the SARS-CoV-2 Nucleocapsid Protein C-Terminal Domain Reveals a Novel Recognition Mechanism for Viral Transcriptional Regulatory Sequences. Front Chem 8, 624765 (2020).

57. M. Takeda et al., Solution structure of the c-terminal dimerization domain of SARS coronavirus nucleocapsid protein solved by the SAIL-NMR method. J Mol Biol 380, 608–622 (2008).

58. R. Zhou, R. Zeng, A. von Brunn, J. Lei, Structural characterization of the C-terminal domain of SARS-CoV-2 nucleocapsid protein. Mol Biomed 1, 2 (2020).

59. M. P. Robertson et al., The structure of a rigorously conserved RNA element within the SARS virus genome. PLoS Biol 3, e5 (2005).

60. E. Nikolakaki et al., RNA association or phosphorylation of the RS domain prevents aggregation of RS domain-containing proteins. Biochim Biophys Acta 1780, 214–225 (2008).

61. C. R. Carlson et al., Phosphoregulation of Phase Separation by the SARS-CoV-2 N Protein Suggests a Biophysical Basis for its Dual Functions. Mol Cell 80, 1092-1103.e1094 (2020).

62. S. Wang et al., Targeting liquid–liquid phase separation of SARS-CoV-2 nucleocapsid protein promotes innate antiviral immunity by elevating MAVS activity. Nature Cell Biology 23, 718–732 (2021).

63. F. Borghese, T. Michiels, The Leader Protein of Cardioviruses Inhibits Stress Granule Assembly. Journal of Virology 85, 9614–9622 (2011).

64. F. Borghese, F. Sorgeloos, T. Cesaro, T. Michiels, The Leader Protein of Theiler’s Virus Prevents the Activation of PKR. J Virol 93 (2019).

65. L. J. Visser et al., Foot-and-Mouth Disease Virus Leader Protease Cleaves G3BP1 and G3BP2 and Inhibits Stress Granule Formation. J Virol 93 (2019).

66. A. Kaminski, G. J. Belsham, R. J. Jackson, Translation of encephalomyocarditis virus RNA: parameters influencing the selection of the internal initiation site. Embo j 13, 1673–1681 (1994).

67. S. Nabeel-Shah et al., SARS-CoV-2 nucleocapsid protein binds host mRNAs and attenuates stress granules to impair host stress response. iScience 25, 103562 (2022).

68. Y. Zheng et al., SARS-CoV-2 NSP5 and N protein counteract the RIG-I signaling pathway by suppressing the formation of stress granules. Signal Transduction and Targeted Therapy 7, 22 (2022).

69. C. A. de Haan, H. Vennema, P. J. Rottier, Assembly of the coronavirus envelope: homotypic interactions between the M proteins. J Virol 74, 4967–4978 (2000).

70. D. C. Dinesh et al., Structural basis of RNA recognition by the SARS-CoV-2 nucleocapsid phosphoprotein. PLoS Pathog 16, e1009100 (2020).

71. K. M. Scherer et al., SARS-CoV-2 nucleocapsid protein adheres to replication organelles before viral assembly at the Golgi/ERGIC and lysosome-mediated egress. Sci Adv 8, eabl4895 (2022).

72. L. G. Thorne et al., Evolution of enhanced innate immune evasion by SARS-CoV-2. Nature 602, 487–495 (2022).

73. S. Leary et al., Generation of a Novel SARS-CoV-2 Sub-genomic RNA Due to the R203K/G204R Variant in Nucleocapsid: Homologous Recombination has Potential to Change SARS-CoV-2 at Both Protein and RNA Level. Pathog Immun 6, 27–49 (2021).

74. S. Lu et al., The SARS-CoV-2 nucleocapsid phosphoprotein forms mutually exclusive condensates with RNA and the membrane-associated M protein. Nature Communications 12, 502 (2021).

75. Z. Yang et al., Engagement of the G3BP2-TRIM25 Interaction by Nucleocapsid Protein Suppresses the Type I Interferon Response in SARS-CoV-2-Infected Cells. Vaccines (Basel) 10 (2022).

76. G. M. Duke, A. C. Palmenberg, Cloning and synthesis of infectious cardiovirus RNAs containing short, discrete poly(C) tracts. J Virol 63, 1822–1826 (1989).

77. K. J. Livak, T. D. Schmittgen, Analysis of relative gene expression data using real-time quantitative PCR and the 2(-Delta Delta C(T)) Method. Methods 25, 402–408 (2001).

