## Supplemental data for "SARS-CoV-2 nucleocapsid protein inhibits the PKR-mediated integrated stress response through RNA-binding domain N2b"

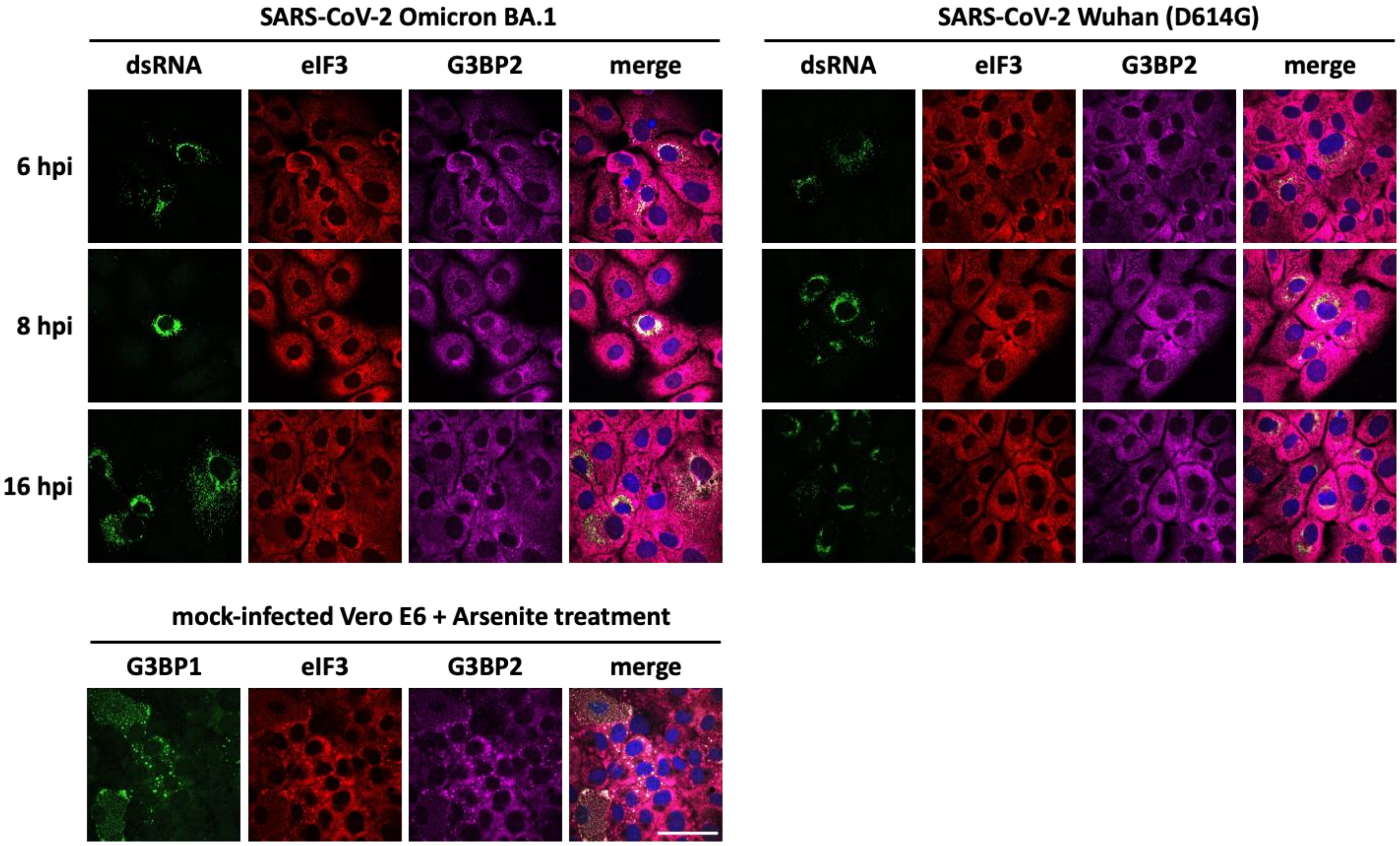

**Fig. S1. SARS-CoV-2 infected Vero E6-TMPRSS2 are virtually devoid of SGs.** Vero E6 cells stabling expressing TMPRSS2 were infected with SARS-CoV-2 Omicron BA.1 and Wuhan (D614G) variants with a MOI of 5. After 6, 8 and 16hpi cells were fixed and stained with antibodies against dsRNA as active replication marker, eIF3 and G3BP2 as SGs markers. Mock-infected cells were treated with Sodium Arsenite to induce SGs and stained with antibodies against G3BP1, G3BP2 and eIF3. Scale bar: 50 µm.

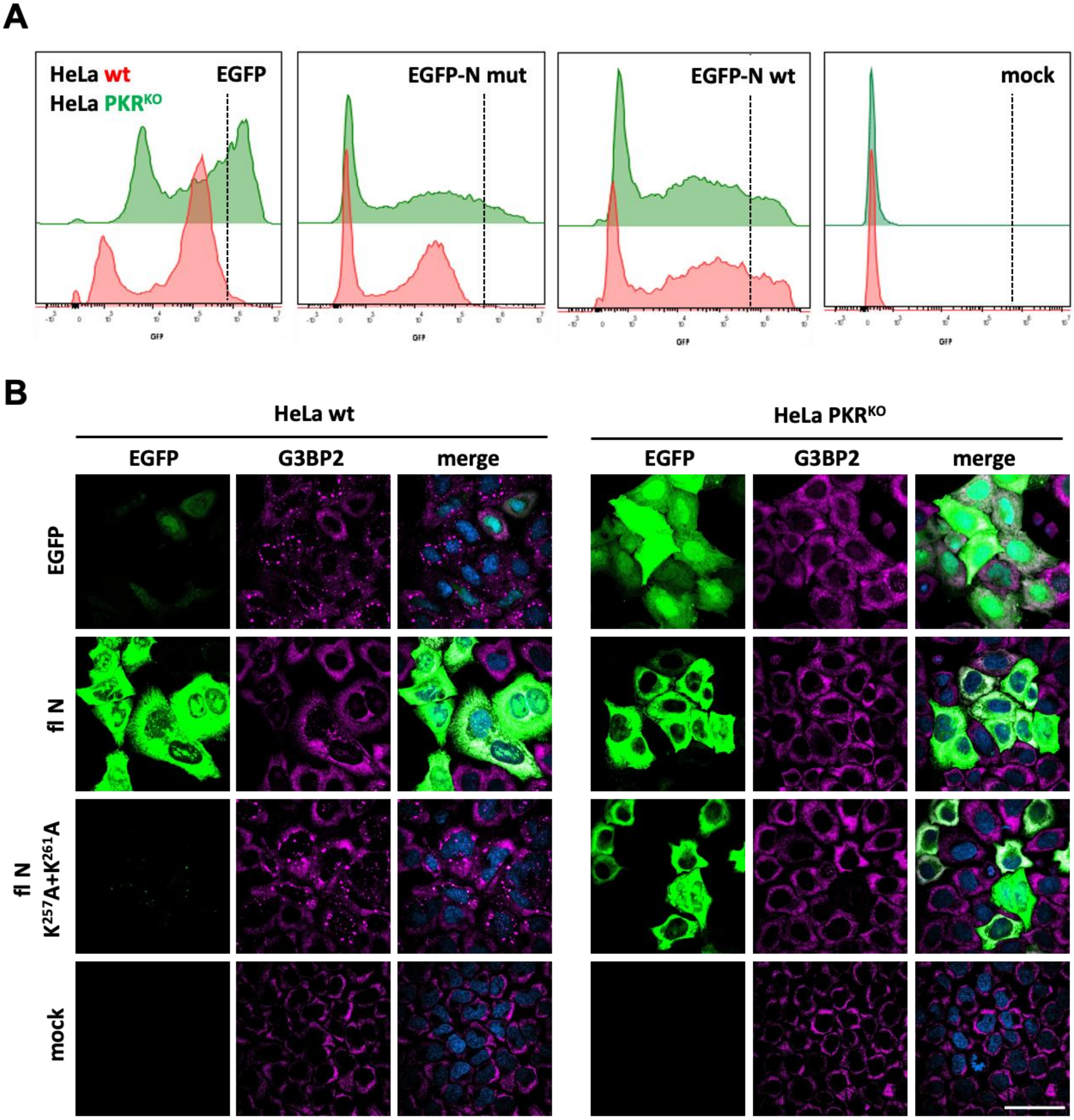

**Fig. S2.** (Related to Fig. 1). **Basal expression levels of EGFP-N and EGFP.** Hela wt and HeLa $PKR^{KO}$ cells were (mock)transfected to express EGFP, EGFP-N and $EGFP\text{-}N^{mut}$, i.e. a derivative unable to inhibit the ISR. Expression levels were assessed by (*A*) flow cytometry or (*B*) fluorescence microscopy as in Fig. 1A. The EGFP-N fusion protein is non-codon optimized and three times larger in size than codon-optimized EGFP. Note that expression of EGFP, as indicated by average fluorescence intensity, is higher than that of EGFP-N when compared in HeLa $PKR^{KO}$ cells, i.e. in the absence of translational arrest. However, also note that in HeLa wt cells, under conditions of PKR-mediated ISR-induced translational arrest, (i) expression of EGFP is restricted and (ii) in result, expression of EGFP-N greatly exceeds that of EGFP in a sizeable population of transfected cells. The analysis has been performed for all the constructs used in this study; data available upon request.

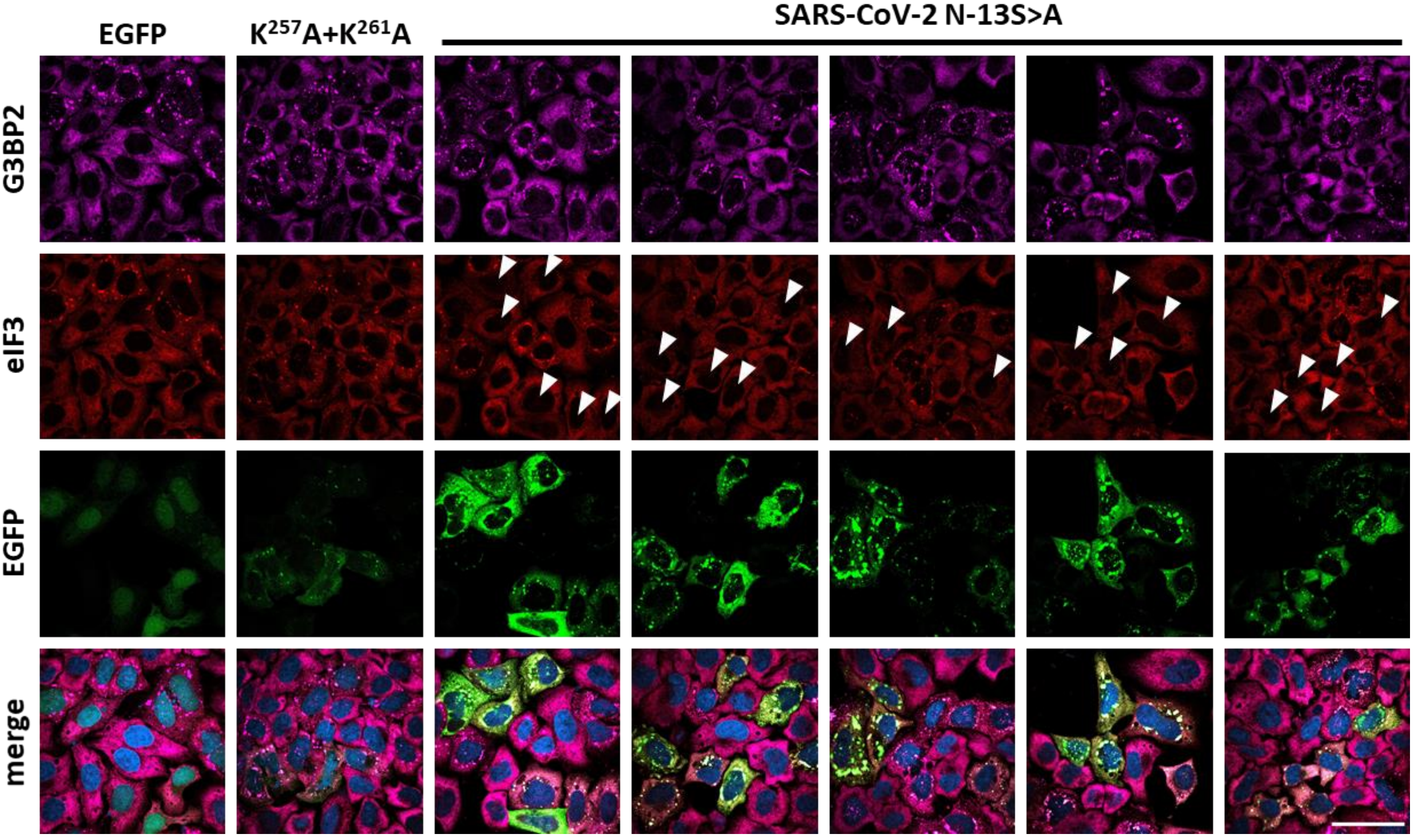

**Fig. S3** (related to Fig.4)**. Phosphorylation of SR motif of SARS-CoV-2 N affects N aggregation but not suppression of the ISR-induced translation arrest and SG formation.** HeLa R19 cells were transfected with SARS-CoV-2 N mutant containing Ala substitutions of 13 Ser (13S>A) in the SR motif to prevent phosphorylation. Transfected cells were stained for eIF3 and G3BP2 as markers for SGs. Most of the transfected cells with 13S>A N mutant show G3BP2 aggregates but not positive for eIF3 (white arrows). Size bar: 50µm.

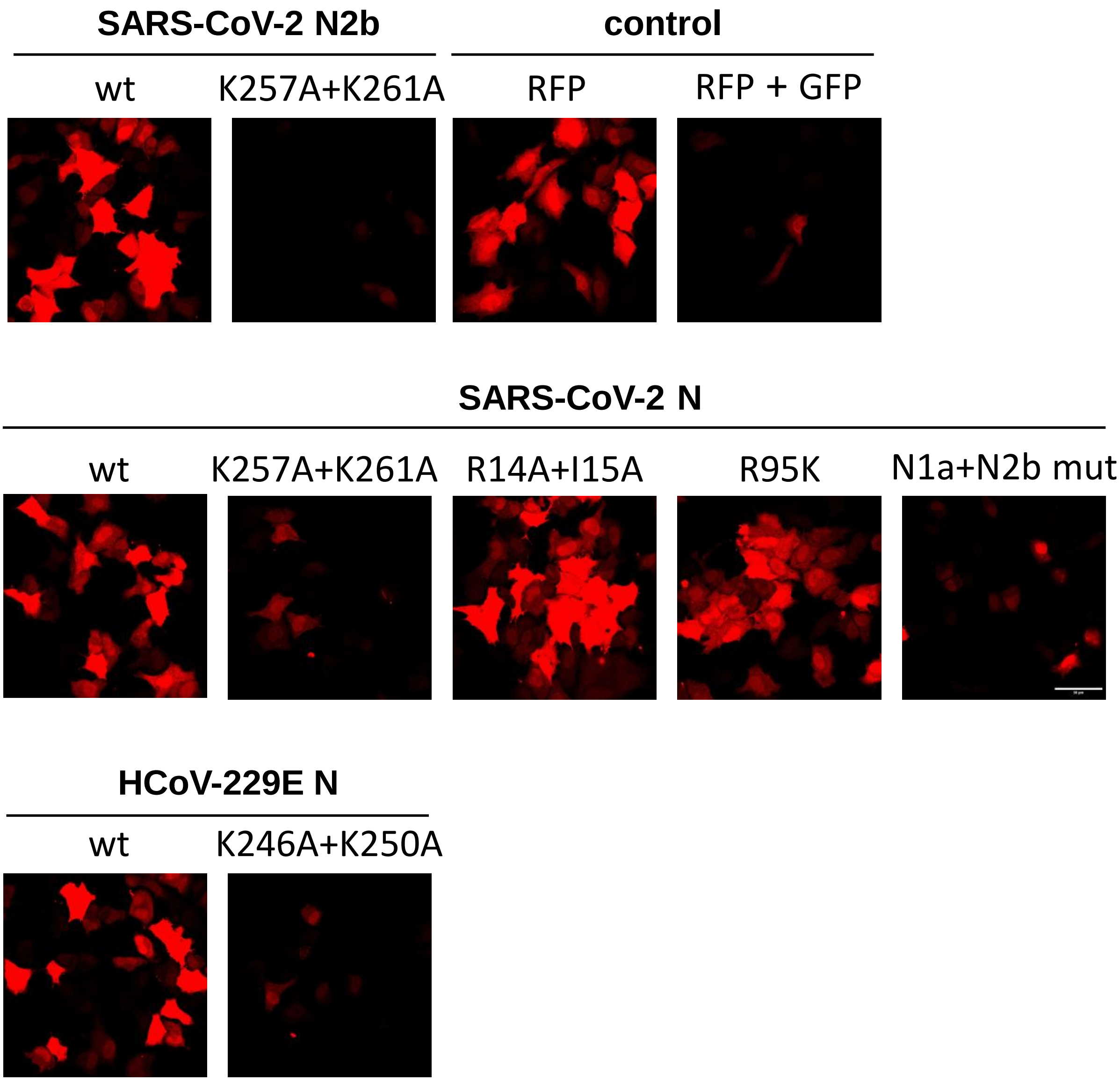

**Fig. S4.** (related to Fig. 6). RFP expression of HeLa cells co-transfected with RFP and indicated EGFP-fused proteins. Scale bar: 50 µm.

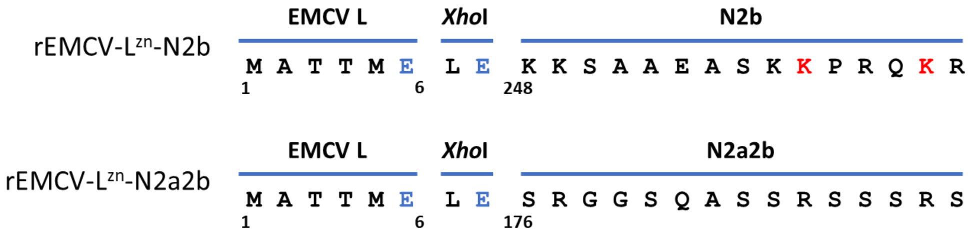

**Fig. S5.** (related to Fig. 7). N-termini of the rEMVC-$L^{zn}$-N2b and rEMVC-$L^{zn}$-N2a2b polyproteins. Indicated are the N-terminal six aa residues of EMVC L, the two residues encoded by an engineered *Xho*I cleavage site fused to SARS CoV-2 N residues 248-365 or 176-365, respectively.

**Fig. S5**

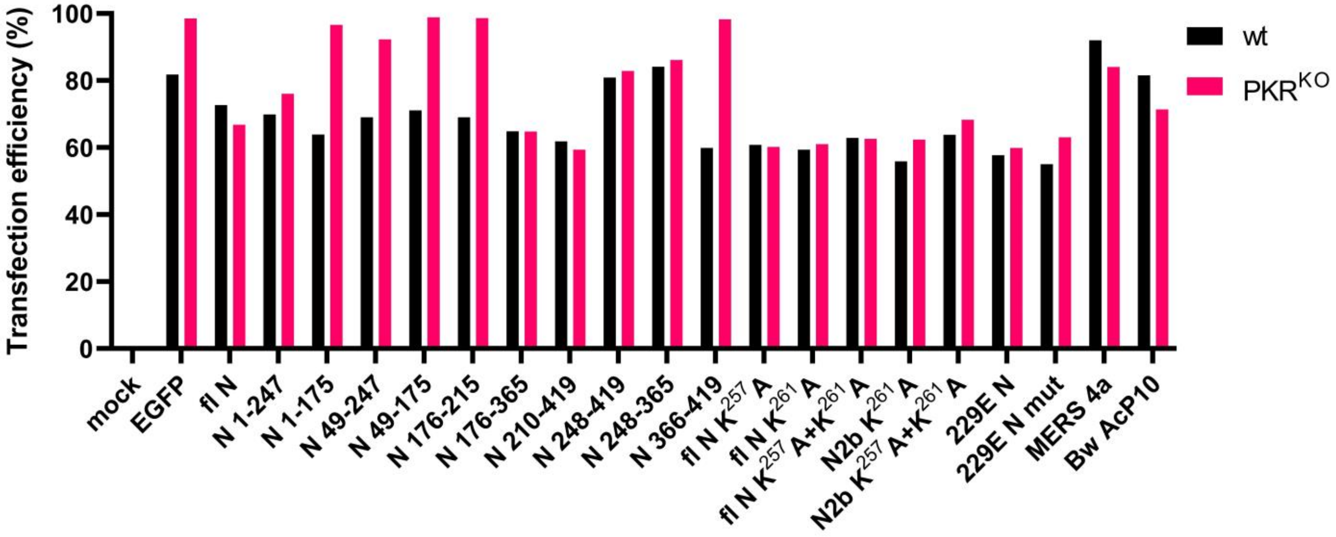

**Fig. S6. Transfection efficiency of N-derivate constructs in HeLa wt and HeLa $PKR^{KO}$ cells.** Transfection efficacy of transfected constructs used throughout this study has been measured by flow-cytometry in HeLa-R19 wt and HeLa $PKR^{KO}$ cells.

**Fig. S6**

| Comparison | Decrease % | Significance | P value |
| --- | --- | --- | --- |
| GFP vs N (1-419 aa) | 95,9 | **** | <0,0001 |
| GFP vs 1-247 aa | none | ns | 0,9998 |
| GFP vs 1-175 aa | none | ns | 0,9508 |
| GFP vs 49-247 aa | none | ns | 0,3596 |
| GFP vs 49-175 aa | none | ns | 0,8692 |
| GFP vs 175-215 aa | none | ns | >0,9999 |
| GFP vs 175-365 aa | 92,8 | **** | <0,0001 |
| GFP vs 175-419 aa | 95,9 | **** | <0,0001 |
| GFP vs 210-419 aa | 97,7 | **** | <0,0001 |
| GFP vs 247-419 aa | 94,1 | **** | <0,0001 |
| GFP vs 247-365 aa | 97,1 | **** | <0,0001 |
| GFP vs 366-419 aa | none | ns | 0,9992 |

**Table S1** (related to Fig. 2A). Comparison of mean percentages of transfected cells with SGs calculated from three biological triplicates, showing the decrease in %, as compared to the control (pEGF-N3-transfected cells, “GFP”). Ordinary One-way ANOVA, Dunnett's multiple comparison test.

### Table S1

| SARS-CoV-2 N2b domain | | | |
|---|---|---|---|
| **Comparison** | **Decrease %** | **Significance** | **P value** |
| GFP vs wt | 97,8 | **** | <0,0001 |
| GFP vs K257A | 58,3 | **** | <0,0001 |
| GFP vs K261A | none | ns | 0,7482 |
| GFP vs K257A+K261A | none | ns | 0,7482 |
| GFP vs Q272A+Q289A | 95 | **** | <0,0001 |
| GFP vs R276A+R293A | 95,5 | **** | <0,0001 |
| **Full-length SARS CoV-2 N** | | | |
| **Comparison** | **Decrease %** | **Significance** | **P value** |
| GFP vs N wt | 96,2 | **** | <0,0001 |
| GFP vs K257A | none | ns | 0,9707 |
| GFP vs K261A | none | ns | 0,8721 |
| GFP vs K257A+K261A | none | ns | 0,4768 |
| GFP vs R276A+R294A | 93,7 | **** | <0,0001 |
| **Full-length HCoV-229E N** | | | |
| **Comparison** | **Decrease %** | **Significance** | **P value** |
| GFP vs wt | 96,4 | **** | <0,0001 |
| GFP vs K246A+K250A | none | ns | 0,3099 |

**Table S2** (related to Fig. 3D). Comparison of mean percentages of transfected cells with SGs calculated from three biological triplicates, showing the decrease in % as compared to the control (pEGF-N3-transfected cells, “GFP”). Ordinary One-way ANOVA, Dunnett's multiple comparison test.

### Table S2

**Total GFP+ cells**

| Comparison | Increase % | Significance | P value |
|---|---|---|---|
| wt vs 1A mut | 24 | **** | <0,0001 |
| wt vs 2A mut | 24 | **** | <0,0001 |
| wt vs 3A mut | 24 | **** | <0,0001 |
| wt vs 4A mut | 24 | **** | <0,0001 |
| wt vs N2b mut | 6 | ns | 0,3114 |
| wt vs N1a+N2b mut | 31 | **** | <0,0001 |
| N1a+N2b mut vs GFP | none | ns | 0,1845 |
| N1a+N2b mut vs N2b mut | 19 | **** | <0,0001 |
| N1a+N2b mut vs 1A mut | none | ns | 0,2417 |
| N1a+N2b mut vs 2A mut | none | ns | 0,2417 |
| N1a+N2b mut vs 3A mut | none | ns | 0,1845 |
| N1a+N2b mut vs 4A mut | none | ns | 0,1845 |

**High GFP+ cells**

| Comparison | Increase % | Significance | P value |
|---|---|---|---|
| wt vs 1A mut | 67 | **** | <0,0001 |
| wt vs 2A mut | 63 | **** | <0,0001 |
| wt vs 3A mut | 66 | **** | <0,0001 |
| wt vs 4A mut | 65 | **** | <0,0001 |
| wt vs N2b mut | 20 | * | 0,0141 |
| wt vs N1a+N2b mut | 83 | **** | <0,0001 |
| N1a+N2b mut vs GFP | - | * | 0,1845 |
| N1a+N2b mut vs N2b mut | 34 | **** | <0,0001 |
| N1a+N2b mut vs 1A mut | none | ns | 0,0872 |
| N1a+N2b mut vs 2A mut | 11 | * | 0,0185 |
| N1a+N2b mut vs 3A mut | none | ns | 0,0680 |
| N1a+N2b mut vs 4A mut | 10 | * | 0,0407 |

**Table S3** (related to Fig. 5B-C). Comparison of mean percentages of transfected cells with SGs calculated from three biological triplicates, showing the decrease in % as compared to wt N or the mutant N1a+N2b N. Ordinary One-way ANOVA, Dunnett's multiple comparison test.

### Table S3
